# Design of immunogens for eliciting antibody responses that may protect against SARS-CoV-2 variants

**DOI:** 10.1101/2022.01.24.477469

**Authors:** Eric Wang, Arup K. Chakraborty

## Abstract

The rise of SARS-CoV-2 variants and the history of outbreaks caused by zoonotic coronaviruses point to the need for next-generation vaccines that confer protection against variant strains. Here, we combined analyses of diverse sequences and structures of coronavirus spikes with data from deep mutational scanning to design SARS-CoV-2 variant antigens containing the most significant mutations that may emerge. We trained a neural network to predict RBD expression and ACE2 binding from sequence, which allowed us to determine that these antigens are stable and bind to ACE2. Thus, they represent viable variants. We then used a computational model of affinity maturation (AM) to study the antibody response to immunization with different combinations of the designed antigens. The results suggest that immunization with a cocktail of the antigens is likely to promote evolution of higher titers of antibodies that target SARS-CoV-2 variants than immunization or infection with the wildtype virus alone. Finally, our analysis of 12 coronaviruses from different genera identified the S2’ cleavage site and fusion peptide as potential pan-coronavirus vaccine targets.

**Author Summary:** SARS-CoV-2 variants have already emerged and future variants may pose greater threats to the efficacy of current vaccines. Rather than using a reactive approach to vaccine development that would lag behind the evolution of the virus, such as updating the sequence in the vaccine with a current variant, we sought to use a proactive approach that predicts some of the mutations that could arise that could evade current immune responses. Then, by including these mutations in a new vaccine antigen, we might be able to protect against those potential variants before they appear. Toward this end, we used various computational methods including sequence analysis and machine learning to design such antigens. We then used simulations of antibody development, and the results suggest that immunization with our designed antigens is likely to result in an antibody response that is better able to target SARS-CoV-2 variants than current vaccines. We also leveraged our sequence analysis to suggest that a particular site on the spike protein could serve as a useful target for a pan-coronavirus vaccine.

## 1 Introduction

Vaccines that protect against severe acute respiratory distress coronavirus 2 (SARS-CoV-2) are highly effective. However, variants that impact vaccine efficacy are emerging. For example, the Beta (B.1.351) variant significantly reduces neutralizing antibody titers (1, 2), and the Delta (B.1.617.2) variant both increases transmission and reduces neutralizing antibody titers (3–6). Although current vaccines still remain effective at preventing severe illness upon infection with these variants (7), their appearance signals that others more capable at evading antibodies elicited by the current vaccines may emerge. Indeed, available data on the Omicron variant suggest that this is true (8, 9). Moreover, other pathogenic coronaviruses could evolve in the future due to zoonosis. Thus, vaccines that protect against potential SARS-CoV-2 variants and those that protect against potential zoonotic coronaviruses would serve as a shield against future outbreaks.

Neutralizing antibodies can prevent infection by binding to a virus’ surface proteins and preventing the virus from entering cells. In samples taken from convalescent COVID-19 donors, the isolated neutralizing antibodies commonly target the spike’s receptor-binding domain (RBD) (10–12), which is located in the S1 domain and is responsible for facilitating viral entry by binding to angiotensin-converting enzyme 2 (ACE2) (13).

In response to antigen (whether from vaccination or natural infection), antibodies are generated by a Darwinian process known as affinity maturation (AM) that occurs in secondary lymphoid organs (14). Activated germline B cells seed structures known as germinal centers (GC), where they undergo multiple cycles of expansion, mutation, and selection based on the binding affinity of their B cell receptor (BCR) to antigen. Through this process, B cells increase their binding affinity to antigen up to 1000-fold or more. B cells can differentiate into plasma cells that secrete antibodies, which are soluble and modified forms of the BCR that inhibit pathogens through neutralization or various other effector functions (15).

Previous computational models of AM have focused on the response to single antigens (16–21) or the development of broadly neutralizing antibodies (bnAbs) for influenza and human immunodeficiency virus (HIV) upon immunization with variant antigens (22–31). The latter studies on highly mutable viruses aimed to study strategies for induction of bnAbs that target the conserved residues of the viral spike.

However, given that SARS-CoV-2 appears to mutate more slowly than influenza and HIV (32, 33), a different strategy is feasible for protection against its variants. Instead of targeting strictly conserved regions, an immunization scheme that generates an appropriate polyclonal response against variable regions could protect against mutant strains. Antibodies targeting the variable class 1 and class 2 RBD epitopes of SARS-CoV-2 tend to be neutralizing, while antibodies that target the conserved class 4 epitope can be non-neutralizing because of their inability to compete with ACE2 binding (34). Thus, targeting variable regions may also be more likely to generate neutralizing antibodies against SARS-CoV-2 variants.

At least two significant questions need to be answered in order to design such a vaccine that can protect against SARS-CoV-2 variants,

1. Which antigens should be used, and in what combination/order should they be administered, to optimally produce the desired antibody response?
2. How does previous infection or immunization with SARS-CoV-2 affect antibody evolution upon administering the chosen antigens?

In this study, we aimed to address these questions pertinent to vaccination that protects against SARS-CoV-2 variants and beyond. We first developed a method of calculating conservation at different sites of the spike protein by analyzing both structural and sequence data. By applying it to sarbecovirus spike proteins and combining the results with previous deep mutational scanning results, we designed 6 antigens that may protect against the most significant RBD escape mutations in SARS-CoV-2. We then determined that these antigens are stable and bind ACE2 using neural networks that we trained on deep mutational scanning data to predict RBD expression and ACE2 binding from sequence, Finally, we used a computational model of AM to study the antibody response to these antigens. Thus, we identified an immunization scheme which may produce higher titers against current and potential variants than immunization with the wildtype (WT) antigen alone. But, such a vaccination scheme produces lower anti-WT titers than the WT vaccine does, illustrating the fact that a strain-specific vaccine is generally the most effective at protecting against a particular strain. However, a vaccine that adequately protects a population when diverse new variants emerge can serve as a shield and provide time for a strain-specific vaccine to be developed.

Our results also highlight factors that impact titers such as previous exposure to WT antigen and the fraction of GC-seeding cells that are memory cells generated during past exposure. By applying the conservation analysis to 12 diverse coronaviruses, we also identify the S2’ cleavage site and fusion peptide as conserved regions, pointing to a potential target for a pan-coronavirus vaccine.

## 2 Methods

### 2.1 Spike conservation

In order to identify conserved and variable residues in coronavirus spike proteins, we leveraged both sequence and structural data. Structural data can be used to determine structural conservation (as described below) in a way that does not strongly depend on the number of insertions or deletions. Inclusion of structural data has a significant impact on which residues are considered conserved, as illustrated in Figure S1. In our method, each residue of the spike was assigned a conservation fraction ranging from 0 to 1, which was the average of the structural conservation fraction and the biochemical conservation fraction. An overview of the method that we use to determine these conservation scores is provided in Figure 1.

**Figure 1.**
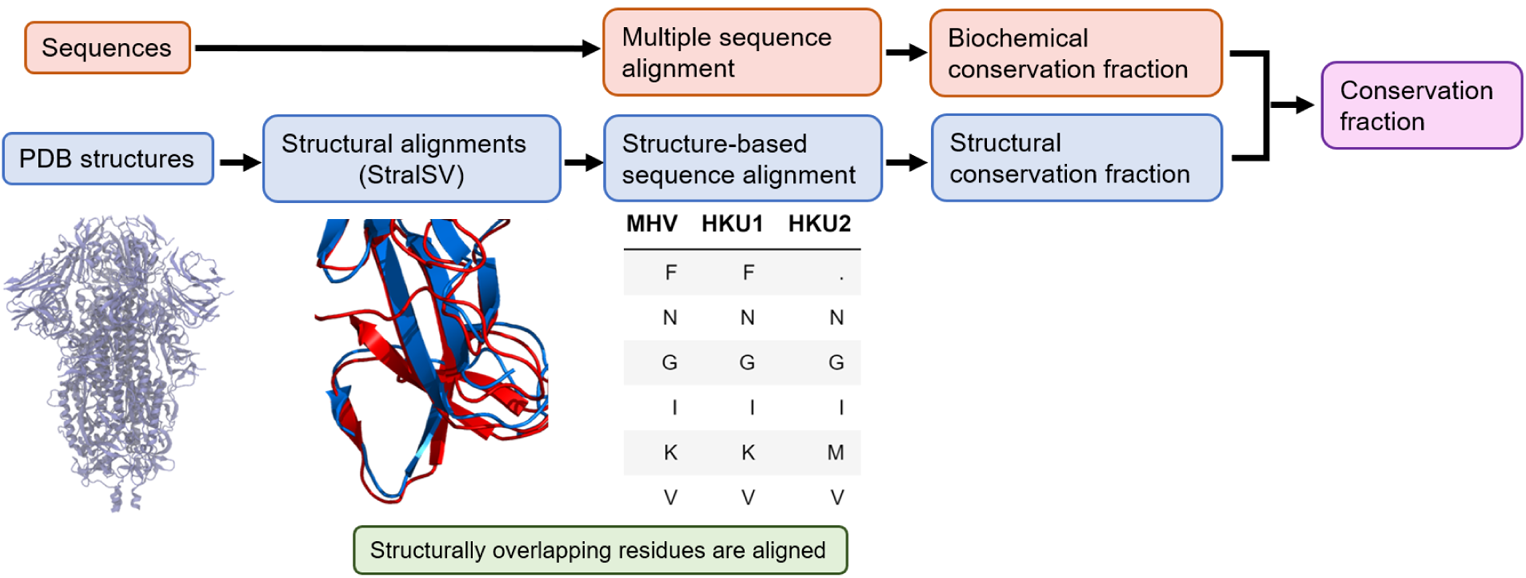
Overview of conservation analysis workflow. Schematic of conservation fraction calculation. Structure analysis (blue) and sequence analysis (red) are independently carried out and averaged to obtain the final conservation fraction.

#### 2.1.1 Structural conservation fraction

Using the structures for the trimeric spike of various coronaviruses (Table S1), atoms were removed from each trimer so that only a monomer remained. Thus, the structural conservation fraction was calculated using a monomer rather than the full trimer, but the monomer retained the conformation that it had in the trimer. The STRAL-SV server (35) was used to align these monomeric spike protein structures. The STRAL-SV server addresses the fact that multi-domain proteins can adopt multiple conformations, such as the spike protein’s RBD being oriented down in one structure and up in another structure, which would have confounded a standard global alignment. In STRAL-SV, a sliding-window approach divided the monomer structures into overlapping fragments of 90 amino acids, and structural alignments were performed between the fragments. This process identified residues across different coronavirus spikes that structurally overlap in the alignment. This data was used to construct a structure-based sequence alignment. A table is first constructed (see Figure 1) where the residues of the spike proteins are the rows, the column names correspond to coronaviruses (such as MHV, HKU1, and HKU2), and the entries of the table are the 20 amino acids or a placeholder entry of “.” The placeholder entry of “.” means that the coronavirus spike protein was not able to align at that residue.

The structural conservation fraction was calculated from the structure-based sequence alignment using a particular coronavirus as a reference. For each residue, the structural conservation fraction was the fraction of other coronaviruses that are able to structurally align at that residue. We illustrate this with an example from Figure 1’s table, assuming for simplicity that the first row corresponds to residue 1. If we select residue 1 and MHV as a reference coronavirus, the other coronaviruses are HKU1 and HKU2. Only HKU1 is able to structurally align, so the structural conservation fraction is 0.5. It is also possible that the reference coronavirus did not align at the chosen residue, in which case the structural conservation fraction is 0. For example, consider residue 1 using HKU2 as a reference coronavirus. Since the entry is “.”, then the structural conservation fraction is 0. Note that Figure 1’s table is truncated and is shown for illustration purposes only.

#### 2.1.2 Biochemical conservation fraction from sequence analysis

For each coronavirus, a set of several hundred spike sequences were collected from the NCBI Protein database and processed (see SI for details of collection and processing). All coronavirus alignments were then concatenated together into a single multiple sequence alignment as illustrated in Figure S2.

The biochemical conservation fraction was calculated from the multiple sequence alignment using a particular coronavirus as a reference. For each residue, the biochemical conservation fraction was the fraction of sequences that have an amino acid in the same class (hydrophobic, polar, positive, or negative) as the reference coronavirus’s consensus (most common) amino acid. Thus, this calculation tolerates mutations if they are biochemically similar.

To illustrate this procedure using Figure S2, assuming that the first row corresponds to residue 1 for simplicity, suppose we wish to calculate the biochemical conservation fraction of residue 1 using MHV as a reference coronavirus. The consensus amino acid from the MHV alignment is F, which is a hydrophobic amino acid. The biochemical conservation fraction is the fraction of amino acids in the first row of the concatenated alignment that are also hydrophobic.

#### 2.1.3 Conservation fraction

The conservation fraction was the average of the structural conservation fraction and the biochemical conservation fraction. For the design of antigens that may protect against SARS-CoV-2 variants, only severe acute respiratory syndrome coronavirus (SARS-CoV) and SARS-CoV-2 data were considered. Since SARS-CoV and SARS-CoV-2 are closely related, the threshold conservation fraction below which a residue was identified as variable was chosen to be 0.99.

For pan-coronavirus vaccine target identification, we considered 12 coronaviruses from different genera (Table S1), which constituted an analysis of 12 spike protein structures and ~4000 spike protein sequences. We set a threshold conservation fraction of 0.8, which was chosen because it was the largest value at which we observed a conserved region comparable in size to an antibody footprint.

### 2.2 Design of cocktail antigens for protection against SARS-CoV-2 variants

We hypothesized that a cocktail composed of variant SARS-CoV-2 spike sequences can generate a polyclonal response that protects against variants of SARS-CoV-2. The aim is to generate a polyclonal response that is composed of strain-specific antibodies that together can protect against multiple variants. We do not aim to elicit bnAbs that target a particular conserved epitope, as in studies being done in the context of universal influenza and HIV vaccines (22–25, 36–41). To design antigens that can elicit the desired polyclonal response, we first generated a list of mutations (with respect to the wildtype Wuhan SARS-CoV-2 sequence) that satisfied the criteria below. The lists of mutations satisfying each of these criteria are provided at https://github.com/ericzwang/sars2-vaccine/tree/main/data/cocktail_design_criteria_mutations.

1. The mutation was in a residue determined to be variable using the calculation outlined in Section 2.1. The structure and sequence data were restricted to SARS-CoV-2 and SARS, and the conservation fraction threshold was set to 0.99. SARS-CoV was included because a calculation exclusively using the structures of SARS-CoV-2 variants and ~300,000 SARS-CoV-2 sequences classified nearly the entire spike protein as conserved (Figure S3). In other words, only a few SARS-CoV-2 mutations have occurred in current SARS-CoV-2 variants, so we consider residues that are different in SARS-CoV as such mutations have the potential to emerge because these viruses are closely related. The Omicron variant emerged after our analyses were completed, but its characteristics support our use of SARS-CoV data. We considered the 15 RBD mutations in the Omicron variant, and found that 12 out of 15 Omicron RBD mutations are classified as variable residues according to our calculation. Additionally, all RBD mutations in previous variants (Alpha, Beta, Delta, Gamma) are considered variable. Therefore, while few SARS-CoV-2 mutations have already emerged, our choice of including SARS-CoV data appears to be predictive of mutations that may emerge in the future.
2. The mutation did not significantly decrease RBD stability. This was determined from deep mutational scanning experiments carried out by Starr et al. (42) that measured the effect of nearly every single RBD mutation on protein expression, a correlate of protein stability. A threshold on the change in expression from the WT (expressed as log_10_(MFI)) was set to −0.2, which was chosen such that mutations in circulating variants would be considered viable. At this stage, multiple-mutation effects were not considered, but they were considered when the final antigens were evaluated using the expression-prediction neural network we developed (described in Section 2.3).
3. The mutation did not significantly decrease ACE2 binding. This was determined from deep mutational scanning experiments (42), which also measured the effects of single mutations on ACE2 binding. The allowed threshold value of the change in affinity from the WT (expressed as log_10_(K*_D,app_*) where K*_D,app_* was in units of M) was set to −0.2, which was chosen such that mutations in circulating variants would be considered viable. As for protein expression, multiple-mutation effects on ACE2 binding were evaluated later using the neural network described in Section 2.3.
4. The mutation significantly abrogated class 1 or class 2 antibody binding. These mutations were found from various deep mutational scanning experiments over 20 class 1/2 antibodies (43–47) (Table S2), in which a mutation’s ability to abrogate binding was quantified as an escape fraction. In the experiments, many cells are generated which express different RBD sequences, and there are multiple cells that express the same RBD. The cells are sorted into an antibody-escape bin based on their inability to bind a fluorescently tagged antibody. The escape fraction is the fraction of cells expressing a particular RBD that are in the antibody-escape bin. An escape fraction of 0 indicates that no cells expressing a particular RBD are in the antibody-escape bin, and a fraction of 1 indicates that all cells expressing that RBD are in the bin. The 34 mutations with the largest mean escape fractions across all antibodies were selected (see SI for explanation for choosing 34 mutations). Among these 34 mutations, some but not all were also present in circulating strains. The absence of some escape mutations was due to the low mutation rate of SARS-CoV-2 and the boost in transmissibility provided by certain non-escape mutations. An example of a prevalent non-escape mutation is N501Y, which first appeared in the Alpha variant and increased ACE2 binding affinity without significantly affecting antibody binding. Although some of the 34 escape mutations have not arisen yet, they may arise in the future as natural and vaccine-induced responses impose selection pressures. So, they were included in our antigens in order to protect against future variants. Indeed, Q493R, one of the most significant class 1/2 escape mutations in the Omicron variant, is found among these 34 escape mutations and has not appeared in previous variants of concern. Note again that Omicron emerged after we completed our analyses.

The procedure outlined above resulted in a set of mutations that were in variable residues and would likely abrogate binding to neutralizing class 1 and 2 antibodies circulating in vaccinated and naturally infected persons, and did not diminish ACE2 binding or decrease spike stability and so would likely be viable viruses. This list of mutations are candidates for inclusion in antigens that may elicit a broadly protective polyclonal antibody response. Additionally, a survey of ~300,000 SARS-CoV-2 sequences from the GISAID database (see SI for details of collection and processing) revealed several mutations prevalent in circulating strains. Among these, 3 mutations (K417T, K417N, and T478K) were the most prevalent mutations that also escape antibodies, but they were not among the 34 mutations with the largest escape fractions from deep mutational scanning. Nonetheless, these 3 mutations were also considered in order to protect against current circulating variants. In total, the list contained 37 mutations split among 10 residues.

Multiple mutations from the list generated using the procedure described above were spatially close together on the RBD. Multiple such mutations on the same RBD was undesirable as it could lead to coupled effects that affect ACE2 binding or RBD stability. Although such effects were later accounted for using the neural networks described in Section 2.3, the data used to train those networks mostly contained RBDs with spatially separated mutations. So, we grouped the mutated residues such that the residues within a group would be maximally separated in space on the RBD. To accomplish this, we divided the 10 mutated residues into all possible combinations of 2 groups of 5 residues, and our goal was to select the combination with the maximum separation between residues. This was accomplished as follows:

1. For a particular combination of two groups, two numbers, *V*_1_ and *V*_2_, were calculated. To determine *V*_1_, consider the 5 residues in the first group positioned on the structure of the RBD. The centers of mass of the 5 residues formed a polygon, and the volume of that polygon was *V*_1_. Similarly, the volume of the polygon formed by the residue centers of mass of the second group was *V*_2_. The polygons were calculated using the ConvexHull utility in SciPy (48).
2. The combination was assigned a score, which was calculated as the minimum of *V*_1_ and *V*_2_ (min(*V*_1_, *V*_2_)).
3. From all possible combinations, the combination with the largest value of min(*V*_1_, *V*_2_) was chosen. Selecting the score as min(*V*_1_, *V*_2_), instead of *V*_1_ + *V*_2_, avoided choosing cases with severe imbalance between the groups, which may have deleteriously impacted the stability of half of our antigens.

For each of the two groups thus selected, we generated 3 sequences with 5 mutations per sequence. Since there are 2 groups, there are 6 sequences total. For each residue, the amino acid mutations were chosen starting from those with the largest escape fractions (thus, those that escape antibodies most). However, the mutations were also chosen such that the amino acids were dissimilar, meaning that a potential mutation of the same class (hydrophobic, polar, positive, negative) as a previously selected mutation was only selected if no dissimilar mutations were available. For example, if the mutation with the largest escape fraction at residue 490 was F490K, then the first sequence used F490K. For the second sequence, the mutation with the second-largest escape fraction at residue 490 was F490R. But, F490R was skipped because R and K are biochemically similar amino acids. The second sequence then moved to the mutation with the third-largest escape fraction, which was F490E, and this mutation was used because E is not biochemically similar to K. This process repeated for the third sequence and then moved on to another residue. After all 5 residues in the first group were thus examined, the process was repeated for the 5 residues in the second group. Since our antigen design method relied on deep mutational scanning experiments that only studied RBD substitutions, insertions or deletions in the RBD and mutations outside the RBD were not considered.

### 2.3 Neural network-based prediction of RBD expression and ACE2 binding affinity

The published data for selecting stable mutations was based on single mutation effects (42), but the designed sequences possessed 5 mutations each. Therefore, it was not known whether the designed sequences were still stable or maintained ACE2 binding. We noted that the deep mutational scanning experiments provided two datasets – one with measured changes in expression compared to WT for multi-mutant RBDs, and another with measured changes in ACE2 binding affinity (data found at https://github.com/jbloomlab/SARS-CoV-2-RBD_DMS/blob/master/results) compared to WT for multi-mutant RBDs. After removing sequences with no mutations or missing labels, the expression dataset contained ~169,000 mutants with 1-7 mutations, and the binding affinity dataset contained ~135,000 mutants with 1-7 mutations.

For the expression dataset, every sequence was one-hot encoded for the 20 amino acids, with unmutated residues represented by a zero vector. This one-hot encoding matrix was element-wise multiplied with a matrix of the single-mutation expression changes to yield a final matrix termed the expression matrix. In one approach, single-mutation effects were assumed to be additive, and the overall change in expression was the sum of the expression matrix. By training a neural network on the dataset, additivity of single-mutations effects did not need to be assumed. The network architecture is illustrated in Figure 2. In this approach, the expression matrix was passed into a 1D convolutional layer (32 filters, kernel size 3, same padding, and ReLu activation), a max pooling layer (pool size of 2, stride of 1, and same padding), a 1D convolutional layer (16 filters, kernel size 3, same padding, and ReLu activation), a max pooling layer (pool size of 2, stride of 1, and same padding), a flatten layer, a dense layer (64 neurons, ReLu activation), another dense layer (32 neurons, ReLu activation), and an output layer of 1 neuron.

**Figure 2.**
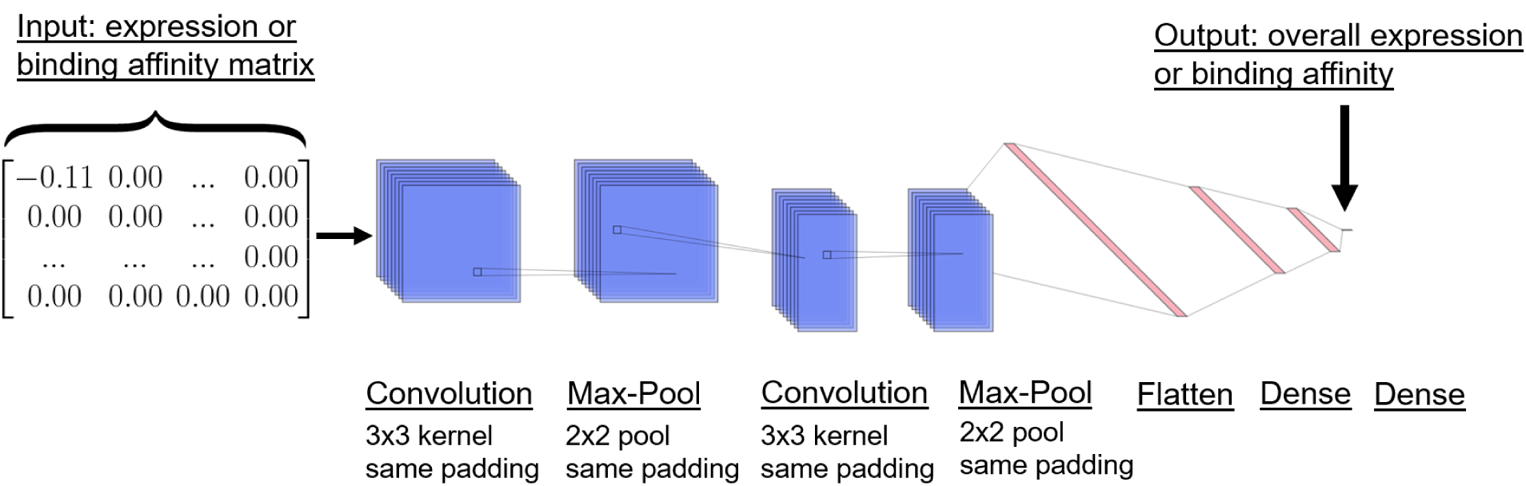
Overview of neural network architecture. Neural network architecture used to predict overall expression or ACE2 binding affinity. The input is either an expression or binding affinity matrix, which is the element-wise product of the one-hot encoded RBD sequence and the matrix of single-mutation changes. The output is the overall expression or binding affinity.

We performed 10-fold cross validation with a 90% train, 5% validation, and 5% test split. Optimization used Adam, a learning rate of 0.001, a batch size of 100, a mean-squared error loss, up to 100 epochs, and early stopping on the validation set with a patience of 3. The model was implemented using Keras version 2.4.0 (49) with the TensorFlow version 2.3.1 backend (50). We calculated the test-set Pearson correlation coefficient on the change in expression.

The same approach was used to train a separate network that predicted changes in ACE2 binding affinity. The only differences were that the binding affinity network was trained using the binding affinity dataset rather than the expression dataset, and the input matrix was the binding affinity matrix instead of the expression matrix. The neural networks were then used to predict the expression and ACE2 binding properties of our designed antigens.

### 2.4 Modeling affinity maturation upon immunization with designed antigens

We simulated the processes that occur during AM using a coarse-grained stochastic model in order to study how antibodies develop in response to immunization with our designed antigens. The goal of these simulations was not to make quantitative predictions, but rather to qualitatively compare different antigen formulations and also understand the underlying mechanisms for the differences. Thus, we aimed to choose the best antigens in a vaccine based on a mechanistic understanding of the GC processes they elicit. An overview of the model is illustrated in Figure 3. This model builds on our past work modeling the affinity maturation response to variant antigens (22–24).

**Figure 3.**
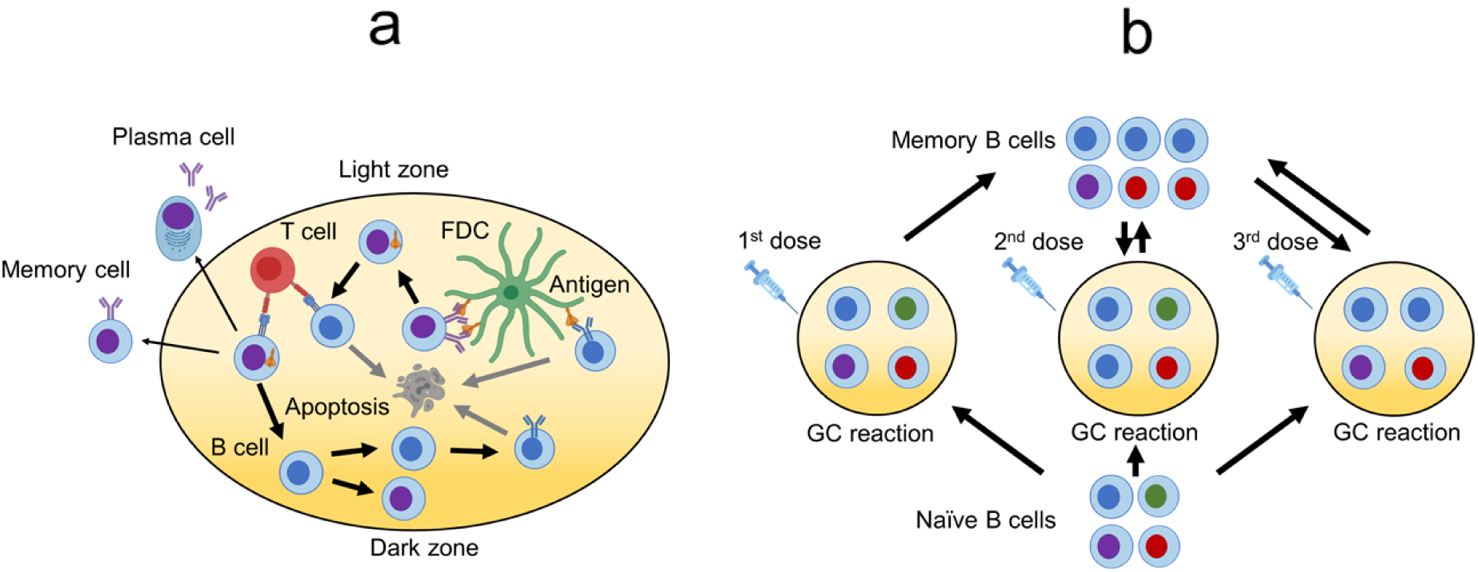
Overview of the affinity maturation model. (a) Schematic of AM within a GC. Activated B cells first undergo expansion and somatic hypermutation (SHM), and then they undergo selection based on their ability to bind and internalize antigen presented on FDCs and receive T cell help. A small fraction of positively selected B cells exits as antibody-secreting plasma cells and memory cells. The others undergo another round of mutation and selection. This cycle repeats until a termination condition is met. (b) Schematic of seeding new GCs upon new immunizations. For the first immunization, the seeding cells come solely from naive B cells. For subsequent immunizations, seeding cells come from both the naive and memory cell populations, and newly generated memory cells join the memory cell population.

#### 2.4.1 Binding free energy representation

The BCR paratope and antigen epitope were each represented by a vector of residues. Both the paratope and epitope were 50 residues long, a length which was chosen based on the number of residues in the class 1 and class 2 epitopes (34, 51–53). For antibodies that did not have the epitope residues reported, the residues were found from the RBD-antibody structure using the PISA server (54–56). This procedure allows identification of RBD residues at the RBD-antibody interface with buried surface areas greater than 0 Å^2^. In order to choose residues that were consistently part of the class 1 or class 2 epitope, only residues that were bound by 3 or more class 1 or class 2 antibodies were included. Using the conservation analysis described above, 9 of the 50 residues had conservation fractions above 0.99 and were assigned as conserved. The remaining 41 residues were assigned as variable.

For a seeding B cell, each residue, k, was associated with a number PAR(*k*), which was initially sampled from a uniform distribution between −0.18 and 0.90. After AM began, the value of PAR(*k*) changed due to mutations, and the bounds on these values were −1 and 1.5. The epitope residues were represented by a number EPI(*k*), with values of +1 for WT residues and negative values for mutated residues depending on how biochemically different the mutated residue is from the WT. For a mutation in the same class (such as hydrophobic to hydrophobic), the epitope residue was −1. For a mutation from hydrophobic to polar, polar to charged, or vice-versa, the epitope residue was −2. For a mutation from hydrophobic to charged or vice-versa, the epitope residue was −3. For a mutation from positively charged to negatively charged or vice-versa, the epitope residue was −4. To illustrate the construction of the epitope vector, we will use the Delta variant, which is defined using the mutations L452R and T478K, as an example. L452R is a hydrophobic to positive mutation (corresponding to a value of −3), and T478K is a polar to positive mutation (corresponding to a value of −2). Therefore, one residue of the epitope vector will have a value of −3, another will have a value of −2, and the remaining residues will have values of +1. The order of the residues does not matter in our simple model.

The binding free energy was calculated according to

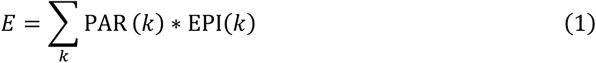

where PAR(*k*) was the paratope residue at position *k* and EPI(*k*) was the epitope residue at position *k*. Essentially, the binding free energy is a dot product between the paratope and epitope vectors. Larger values of *E* corresponded to stronger binding. This coarse-grained representation of the binding free energy was not based on structural considerations and had limitations: (1) the contribution of each residue to the binding free energy was weighted equally; (2) the order of the residues did not matter; (3) only RBD substitutions could be considered; (4) biochemically similar mutations were treated equivalently. The first limitation, that each residue is weighted equally, is a particularly significant assumption since it is known that certain residues more easily abrogate binding upon mutation than others (46). The effect on our model is that our estimates of the potency of antibody responses to variants are likely to be conservative, as weighting all residues equally enables the virus to evade the response more easily as mutations at all sites, rather than a few, can abrogate binding. Additionally, while it is possible to incorporate experimental data in order to reweight the residues, such an approach is not appropriate. This is because the experimental data was measured using antibodies generated from WT infection, but our vaccine will generate different antibodies corresponding to different escape residues. For this reason, we believe that our approach of equally weighting residues is less biased.

That said, we emphasize that this limited representation of the binding free energy is not quantitative and is not meant to accurately reproduce experimental values. Rather, it is a *qualitative* tool meant to facilitate comparison between different antigens and obtain mechanistic insights into AM when used with the rest of the model. The utility of this approach has been demonstrated in previous work, which used a very similar model to provide insights into antibody evolution against HIV antigens that were then validated in animal models (22, 39, 57).

#### 2.4.2 B cell expansion and mutation in the dark zone

Upon initial immunization, 10 naive B cells seeded a new GC. Increasing the number of seeding cells does not change our qualitative results (Figure S4). The B cells then expanded without mutation or selection until the population reached 5120 cells. In AM, activation-induced cytidine deaminase (AID) introduces somatic hypermutations (SHMs) into the BCR. This was modeled by mutating the paratope residues at a rate of 0.14 per sequence per division (B cells divided twice per GC cycle). SHMs had a 0.5 probability of being lethal, 0.3 probability of being silent, and 0.2 probability of modifying the binding energy (58). For mutations that modified the binding free energy, a random residue on the paratope was chosen, and the change in binding free energy for that residue was sampled from a shifted log-normal distribution:

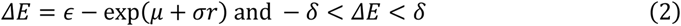

Where *r* was a standard normal random variable, *∊* was a shift parameter, μ was the mean of the log-normal distribution, σ was the standard deviation of the log-normal distribution, and δ limited the maximum Δ*E*. We used the same parameters as Sprenger et. al (∊ = 3, μ = 1.9, σ = 0.5, δ = 1), which were set to match experimental distributions of changes in binding free energies between proteins due to single-residue mutations (59).

#### 2.4.3 Selection in the light zone

In the light zone of the GC, B cells underwent selection based on their binding affinity to antigen. First, each B cell had a probability of successfully internalizing antigen according to

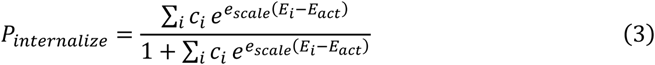

where *i* was a sum over all antigens that the B cell encounters, *c_i_* was the concentration of antigen *i, E_i_* was the binding free energy to antigen *i, E_act_* was a reference free energy, and *e_scale_* was a scaling parameter. We used the same parameters as Sprenger et. al (*e_scale_* = 0.08, *E_act_* = 9*k_b_T*). For a particular immunization scheme, the concentration that produces the highest antibody titers was chosen. The sum in Eq. 3 did not apply for WT doses that only contain a single antigen. For cocktails, *P_internalize_* depended on whether the B cell encounters one antigen at a time or all antigens on the FDC. If the B cell encountered one antigen at a time in each cycle, then a random antigen was chosen, and there was no summation. If the B cell encountered all antigens during each cycle, then the sum included all antigens *i*. In any given cycle, a B cell has a few chances to be positively selected. So, if the antigen concentration on FDCs is sufficiently high, then the sum over all antigens is likely to be more realistic.

B cells that did not internalize antigen underwent apoptosis and were removed from the population, while B cells that internalized antigen competed for T cell help. To model T cell selection if B cells encountered one antigen per cycle, the B cells were ranked according to their binding free energy to the encountered antigen in the last cycle, and the top *F_cut,help_* fraction were positively selected. If B cells instead encountered all antigens per cycle, then the binding free energy ranking was based on all B cell – antigen pairs. The top *F_cut,help_* fraction of pairs were selected, and each B cell survived with a probability equal to the frequency that it appeared in the top *F_cut,help_* pairs. For example, suppose there are 6 antigens and thus 6 B cell – antigen pairs for a particular B cell. If there are 5000 B cells, then there are 30,000 possible B cell – antigen pairs in total. If 4 of the B cell – antigen pairs are in the top *F_cut,help_* ∗ 30,000 pairs, then that B cell survives with a probability of 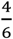. Following Sprenger et. al, we set *F_cut,help_* = 0.7.

#### 2.4.4 Recycling, exit, GC reaction termination, and seeding a new GC

Following Sprenger et. al, positively selected B cells had a 0.7 probability of being recycled for subsequent cycles and a 0.3 probability of exiting the GC as either a plasma or memory cell. The GC reaction stopped if the B cell population exceeded the initial population of 5120 cells, if the number of cycles exceeded 250, or if all B cells died. The first condition is a proxy for the B cells internalizing all the antigen, the second is a proxy for antigen decay over time, and the third is GC extinction. Sprenger et al. set these parameters, along with the parameters from Section 2.4.3 and the bounds on paratope residues from Section 2.4.1, so that a simulation of single-antigen immunization matches experimentally measured GC dynamics of single-antigen immunization (60). Upon a second immunization, a new GC was seeded using 10 B cells, which were a mixture of memory and naive cells. The fraction of memory cells was 1 (all memory cells), except in Section 3.2.4 where we studied the effect of varying this fraction.

#### 2.4.5 Titer calculation

The B cell population at the end of each GC reaction was analyzed for antibody titers. Identical B cells were grouped together as clones, and the binding free energy against an antigen sequence was calculated for each clone. The antibody titer was the total number of B cells that bind to the antigen with a free energy above *E_th_* = 17 *k_b_T*. While the titers quantitatively depended on *E_th_*, the rank order of titers for different immunization schemes that we studied did not. So, the qualitative results did not depend on *E_th_* (Figure S5). The simulation was carried out *N* = 200 times, and the average antibody titer for antigen *i* was calculated as

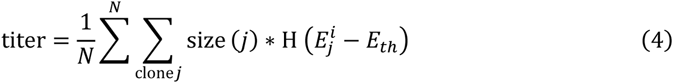

where size (*j*)was the number of B cells belonging to clone *j*, and H represents the step function. Error bars were standard deviations across 10 sets of *N* simulations. The titers thus obtained may be considered to represent the titers generated on average in a typical vaccinated person. This model only considered titers upon completion of each GC reaction. In reality, the titers would be defined by the plasma cells exiting the GC secreting antibodies continuously during the GC reaction.

#### 2.4.6 Mean panel titer calculation

To analyze the coverage provided by antibodies against multiple different antigens, the binding free energy of each B cell clone with a test panel of 100 antigens was calculated. Increasing the panel size to 1000 antigens did not change the results (Figure S6). The antibody titer against each panel antigen was calculated, and the mean panel titer was the mean titer across all 100 panel antigens.

Mean panel titers for several antigen panels were calculated. In the most general panel, residues took values of either +1 or −1, and the probability that the residue is +1 was the conservation fraction. This panel produced almost always resulted in mean panel titers of 0 because the panel antigens had too many mutations at the same time, which are unlikely to emerge in a variant antigen. So, we instead considered panels in which the antigens are restricted to *N_mut_* mutations and varied *N_mut_*. We considered cases in which the *N_mut_* mutations occur in any variable residue and in which the mutations only occur in the 10 residues that are also mutated in the designed antigens. We also considered panels in which mutated residues take on different values (−1, −2, −3, and −4).

## 3 Results

### 3.1 Designed antigens are stable and maintain ACE2 binding based on neural network predictions

Using our conservation analysis on SARS-CoV and SARS-CoV-2 spike proteins and applying the process described in Section 2.2, we obtain the final antigens shown in Table 1. When we trained a neural network on a dataset of ~169,000 mutant RBDs with measured expressions, we obtained a correlation of 0.94±0.01 between measured and calculated values. If the overall change in expression is assumed to be the sum of the single mutation effects, the correlation is instead 0.79±0.01. A separate network trained on a dataset of ~135,000 mutant RBDs to predict ACE2 binding affinity achieves a correlation of 0.97±0.01, compared to 0.78±0.01 for the additive assumption. Thus, while the additive assumption is correlated to the overall change in expression or binding, using network models offers a significant benefit.

**Table 1.** RBD substitutions, changes in expression, and changes in ACE2 binding affinity (calculated using the neural networks) for designed antigens and circulating variants.

| Antigen | RBD substitutions | Change in expression from WT | Change in ACE2 binding affinity from WT |
| --- | --- | --- | --- |
| Sequence 1 | K417F V483R F486P<br>F490Q S494K | $-0.12 \pm 0.04$ | $-0.28 \pm 0.07$ |
| Sequence 2 | K417H V483K F486P<br>F490E S494R | $-0.09 \pm 0.05$ | $-0.53 \pm 0.14$ |
| Sequence 3 | K417Y V483E F486P<br>F490K S494P | $-0.07 \pm 0.05$ | $-0.18 \pm 0.05$ |
| Sequence 4 | L452R T478K E484N<br>Q493K N501A | $-0.06 \pm 0.03$ | $+0.01 \pm 0.27$ |
| Sequence 5 | L452W T478K E484R<br>Q493R N501A | $-0.13 \pm 0.03$ | $-0.20 \pm 0.07$ |
| Sequence 6 | L452E T478K E484K<br>Q493R N501A | $-0.11 \pm 0.03$ | $-0.15 \pm 0.06$ |
| Alpha Variant | N501Y | $-0.11 \pm 0.02$ | $-0.18 \pm 0.06$ |
| Beta Variant | K417N E484K N501Y | $-0.08 \pm 0.02$ | $-0.36 \pm 0.08$ |
| Delta Variant | L452R T478K | $-0.01 \pm 0.02$ | $+0.21 \pm 0.23$ |
| Gamma Variant | K417T E484K N501Y | $-0.06 \pm 0.04$ | $-0.42 \pm 0.12$ |

Since these neural networks achieve high correlations on test sets, we compared the predicted expressions and binding affinities for our designed antigens to several circulating variants (Table 1). None of the designed antigens significantly decrease expression or ACE2 binding affinity more than the corresponding most deleterious circulating variants (Alpha for expression and Gamma for ACE2 binding). Since these circulating variants produce viable virions, we conclude that, upon immunization, our designed antigens would generate immune responses that were relevant for viable viral mutants that may emerge in future. Indeed, although our antigens were designed prior to the appearance of the Omicron variant, mutations in residues 417, 478, 484, 493, and 501 are shared between our designed antigens and the Omicron variant. Residue 493 is particularly notable because it was not mutated in previous variants and is one of the most significant escape residues for evading class 1 and 2 antibodies.

### 3.2 Modeling the affinity maturation response to antigens

We then used our computational model of AM to study how antibodies develop in response to our antigens as well as the WT spike protein. The goals were to identify vaccination schemes using our antigens that may be best at protecting against variants compared to vaccination with the WT spike alone, and to get mechanistic insights into the pertinent AM processes.

#### 3.2.1 Reduced titers against SARS-CoV-2 variants following WT immunization

We simulate AM following one WT immunization, which models infection or one vaccine dose, and two WT immunizations, which models two vaccine doses or infection followed by one vaccine dose. In Figure 4a, the anti-WT titers are high for one immunization (blue bars in various panels) and even higher for two immunizations (orange bars in various panels). In contrast, the titers against variants are reduced depending on the mutational distance between the variant and WT. For example, following two WT immunizations, anti-Beta/Gamma titers are reduced ~6-fold relative to anti-WT titers, anti-Delta titers are reduced ~2.5-fold, and anti-Alpha titers are nearly equivalent to anti-WT titers. This is due to the mutational distances of these variants as the Beta and Gamma variants have 3 RBD substitutions, the Delta variant has 2 RBD substitutions, and the Alpha variant has 1 RBD substitution. This also recapitulates experimental data demonstrating that the Alpha variant minimally reduces neutralizing titers while Beta, Delta, and Gamma significantly reduce titers (1, 2, 4–6, 61–65). These studies report differences between anti-Beta and anti-Gamma titers due to the importance of specific mutations or mutations outside of the RBD, but this coarse-grained model is not structurally detailed enough to recapitulate those differences.

**Figure 4.**
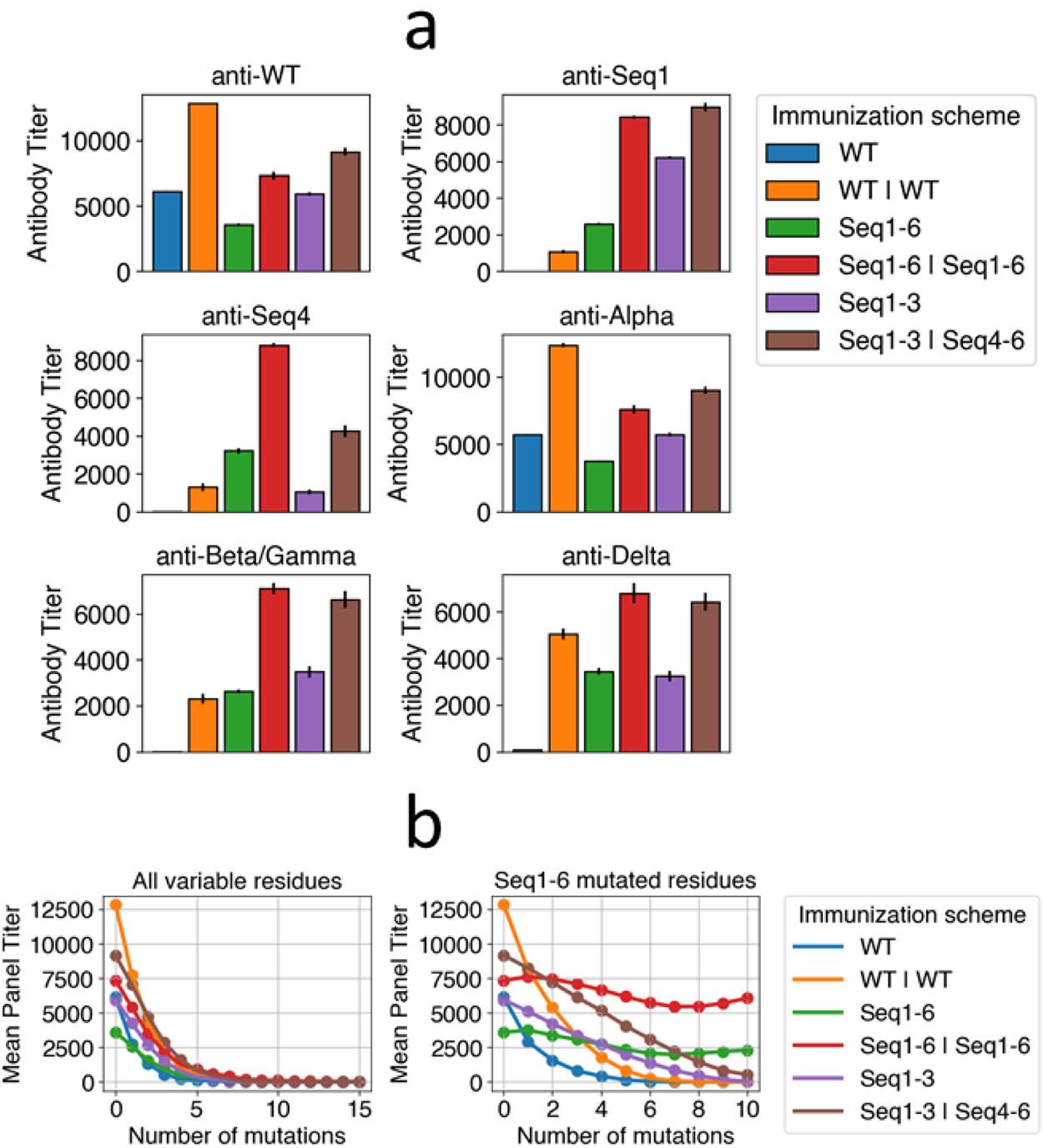
Titers against certain variants are low following WT immunization but high following cocktail immunizations with designed antigens. (a) Titers against WT and variants following different WT or cocktail immunization schemes. Sequences 2 and 3 are similar to sequence 1, so they are omitted for clarity. Sequences 5 and 6 are also omitted because they are similar to sequence 4. Beta and Gamma variants carry biochemically similar RBD mutations, so they are equivalent in this model. (b) Mean panel titers as a function of the number of mutations in panel antigens for different immunization schemes. Two types of panels are considered: one in which panel antigens can be mutated in any variable residue (All variable residues), and one in which panel antigen mutations can only occur in the 10 residues that are also mutated in sequences 1-6 (Seq1-6 mutated residues). It is possible for the panel antigens to have more mutations than in sequences 1-6, which have 5 mutations each. For example, for n=8, every panel antigen has 8 mutations among the 10 residues that were mutated in the designed antigens.

#### 3.2.2 Titers are higher for immunization with a cocktail of designed antigens than WT and depend on the choice of antigens in the cocktail

We then used our model to identify an optimal immunization scheme with our 6 designed antigens. We simulated a single immunization with all 6 antigens (Seq1-6), two immunizations with all 6 antigens (Seq1-6 | Seq1-6), a single immunization with sequences 1-3 (Seq1-3) and an immunization of sequences 1-3 followed by an immunization with sequences 4-6 (Seq1-3 | Seq4-6). We assume that the antigens are homogeneously distributed on the FDCs at high concentrations, so a B cell encounters all antigens in each cycle, but assuming that a B cell encounters one antigen in each cycle does not affect the results (see SI for details). The titers are reported in Figure 4a using different colored bars.

For anti-WT titers, WT immunizations provide greater protection than cocktail immunizations, which is expected because the cocktail antigens are mutated from the WT. However, the boosted cocktails (Seq1-6 | Seq1-6 and Seq1-3 | Seq4-6) still retain moderately high titers against WT, indicating that they may still be protective. The immunizations that start with Seq1-3 (Seq1-3 and Seq1-3 | Seq4-6) have slightly higher anti-WT titers than the analogous immunization starting with Seq1-6 (Seq1-6 and Seq1-6 | Seq1-6). This is because the antigens in Seq1-6 are mutated across 10 residues while the antigens in Seq1-3 are mutated across 5 residues, so the antibodies resulting from immunization with Seq1-3 will have more residues that resemble the WT sequence (more positive in the model). Since more positive residues increases the binding free energy with WT antigen, immunization with Seq1-3 results in higher anti-WT titers. The Alpha variant is similar to the WT, so the same reasoning applies to anti-Alpha titers.

Anti-Seq1 titers are low following WT immunization because sequence 1 contains numerous mutations. Seq1-6 immunization produces lower anti-Seq1 titers than Seq1-3 immunization because sequences 2 and 3 contain mutations in the same residues as sequence 1 while sequences 4-6 contain mutations in different residues from sequence 1. However, immunization with Seq1-6 | Seq1-6 versus Seq1-3 | Seq4-6 are nearly equivalent in anti-Seq1 titers. This is because the anti-Seq1 antibodies provided by the boost with Seq1-6 are much greater than the anti-Seq1 antibodies provided by the boost with Seq4-6, which compensates for the lower anti-Seq1 antibodies generated by priming with Seq 1-6 compared to Seq 1-3.

Anti-Seq2 and anti-Seq3 titers showed similar results to anti-Seq1 titers, so they are omitted for brevity. These titers are similar, even though sequence 2 and 3 have different mutations from sequence 1, because the mutations are in the same residues. Paranthetically, this phenomenon of titers mostly depending the site of mutation rather than the actual mutation is observed to some extent in experimental data, as escape residues often can abrogate antibody binding through numerous possible mutations (44), but there are some cases of biochemically similar mutations differentially affecting antibody binding. Our finding noted above illustrates the qualitative, but not quantitative, nature of our model.

For anti-Seq4 titers, immunization with Seq1-6 produces higher titers than immunization with Seq1-3 because sequences 1-3 are all mutated in different residues from sequence 4. Immunization with Seq1-6 | Seq1-6 produces higher anti-Seq4 titers than immunization with Seq1-3 | Seq4-6, which is somewhat unexpected because one might expect that a boost with Seq4-6 would produce higher anti-Seq4 titers than a boost with Seq1-6. The origin of this result is that, because Seq 4-6 are very different from Seq 1-3, the memory B cells produced by priming with Seq 1-3 are poorly adapted to Seq 4-6. Therefore, the boost with Seq4-6 results in many memory B cells in the secondary GC dying (Figure S7), which reduces the anti-Seq4 titer from the boost. Similar to how anti-Seq2 and anti-Seq3 titers are similar to anti-Seq1 titers, anti-Seq5 and anti-Seq6 titers are similar to anti-Seq4 titers and are not shown for brevity.

For anti-Beta/Gamma and anti-Delta titers, cocktail immunization universally produces higher titers than WT immunization with the same number of immunizations (e.g. Seq1-6 is higher than WT and Seq1-6 | Seq1-6 is higher than WT | WT). The titers resulting from immunizations with different cocktails are roughly equivalent, such as the titers resulting from Seq1-6 | Seq1-6 versus Seq1-3 | Seq4-6.

Figure 4b shows the mean panel titer as a function of the number of mutations in the panel antigens from immunization with WT and cocktails. Mutated residues change from 1 to −4, which is chosen because the vaccine should ideally protect against emergent strains with biochemically dissimilar mutations. Using values such as −3, −2, or −1 instead of −4 does not change the qualitative results (Figure S8). If the panel mutations occur in any variable residue (Figure 4b, left panel), then the mean panel titer decreases at a similar rate for the cocktails as for the WT immunization. This is because there are few conserved residues, so the contribution of conserved regions to the binding free energy is not sufficient to generate bnAbs that can protect against strains bearing many mutations at other residues. This is consistent with experiments showing that few antibodies targeting class 1 or class 2 epitopes of the RBD are broadly protective against all single RBD substitutions (46) let alone those bearing multiple such mutations.

We next studied whether immunization with our designed antigens can protect against panels of variants with mutations only in the residues that are also mutated in the designed antigens (Figure 4b, right panel). Note that the actual amino acids at the 10 residues can be different from those in our designed antigens. In this case, the mean panel titers do not significantly drop for immunization with Seq1-6 and Seq1-6 | Seq1-6. That is, vaccination using these schemes protects against variants with up to 10 mutations in particular variable residues. The mean panel titers for Seq1-3 | Seq4-6 do drop as the number of mutations increases, but less so than WT immunization. These results suggest that Seq1-6 | Seq1-6 may be an effective immunization scheme that could protect against current variants and some that may emerge in the future.

#### 3.2.3 Titers against variants depend on previous exposure to WT and the number of mutations in the variant

Many individuals have been previously exposed to WT SARS-CoV-2 either through vaccination or natural infection, which might affect the antibody response generated upon immunization with a cocktail of our designed antigens. This is because memory B cells generated during prior exposure may compete with naïve B cells that seed GCs. To study this process, we use our model to predict antibody response given previous WT immunizations. Figure 5 graphs antibody titer as a function of the number of previous immunizations with WT for different immunization schemes. Although naive cells are likely involved in seeding new GCs *in vivo* (66), here we assume that only memory cells seed new GCs to maximally illustrate the effects of memory. Section 3.2.4 studies the effect of including naive cells.

**Figure 5.**
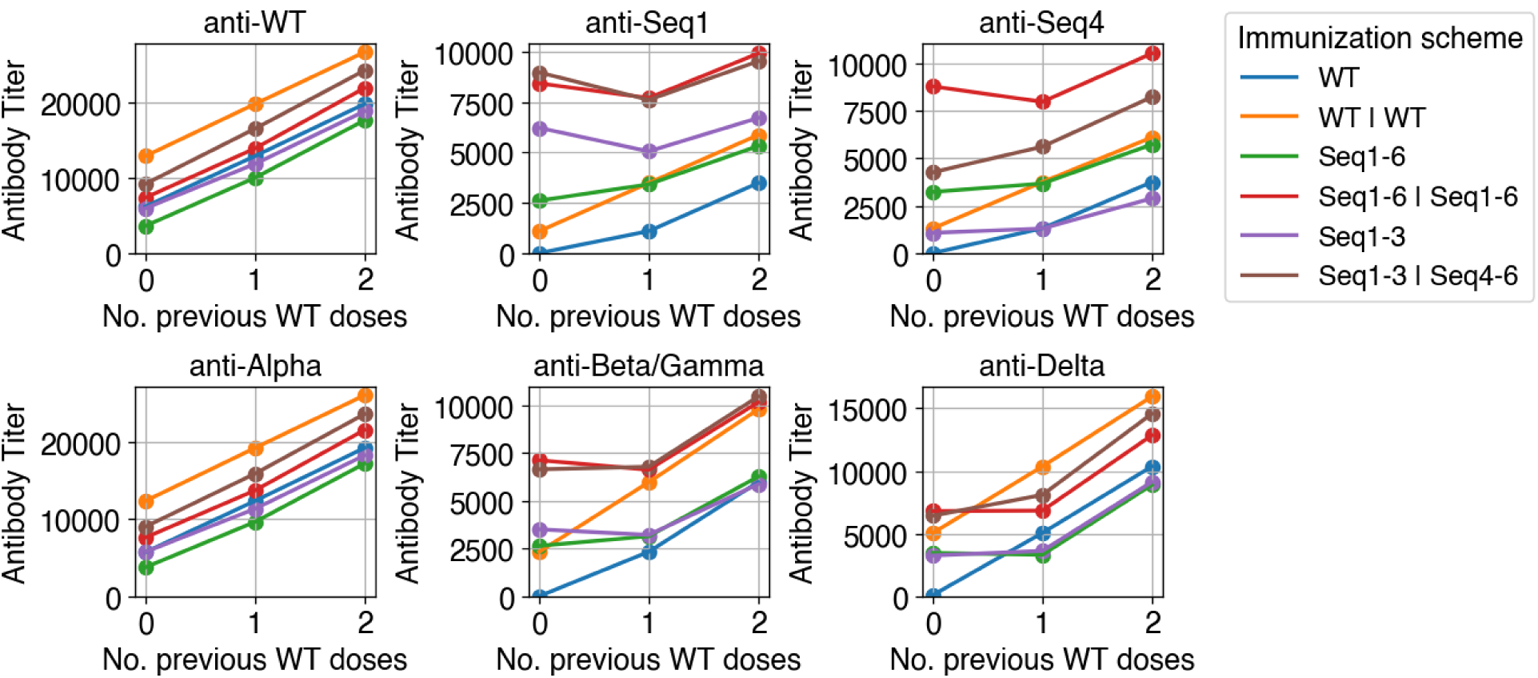
Titers against variants are affected by previous WT immunizations. Titers against various antigens (indicated in panel title) as a function of the number of previous WT immunizations. Sequences 2 and 3 are similar to sequence 1, so they are omitted for clarity. Sequences 5 and 6 are also omitted because they are similar to sequence 4. Beta and Gamma variants carry biochemically similar RBD mutations, so they are equivalent in this model.

First, we compare WT versus WT | WT and WT | WT | WT immunization as well as cocktail versus WT | cocktail and WT | WT | cocktail immunization. Intuitively, anti-WT and anti-Alpha titers steadily increase as the number of previous immunizations with WT increase. However, titers against variants with numerous mutations are not always higher for WT | cocktail and WT | WT | cocktail compared to just cocktail immunization. For example, the anti-Seq1 titers decrease from 0 to 1 previous WT immunization for Seq1-6 | Seq1-6 and Seq1-3 | Seq4-6. This is because the mutational distance between the previous WT immunizations and the cocktails induces GC extinction, thereby limiting antibody development. Nonetheless, the decrease is not large even though GCs are seeded entirely by memory cells.

Next, we compare WT followed by WT immunization versus WT followed by cocktail immunization (for example, WT | WT versus WT | cocktail). For variants with numerous mutations, immunization with the cocktails after the first exposure to WT show higher titers than boosting with WT. For example, anti-Seq1 and anti-Seq4 titers are significantly higher in the WT | Seq1-6 | Seq1-6 and WT | Seq1-3 | Seq4-6 immunizations than the WT | WT immunization. This is because Seq1 and Seq4 each have numerous mutations, so the WT immunizations are not protective against Seq1 and Seq4. Anti-Delta, anti-Alpha, and anti-WT titers are higher for WT | WT immunization than WT | cocktail immunization because of the low number of mutations in these variants. Since the Beta and Gamma variants have more mutations than the Delta and Alpha variants, the anti-Beta/Gamma titers are equivalent between WT | cocktail and WT | WT immunizations.

#### 3.2.4 Sequential immunization of WT and cocktail produces significantly higher titers against current variants than sequential immunization of WT and WT when naive B cells also seed new GCs

Anti-Beta/Gamma and anti-Delta titers are not higher for cocktail immunizations compared to WT immunizations under specific conditions (assuming 1-2 previous immunizations with WT have been given and new GCs are seeded entirely by memory cells). However, if naive cells also seed GCs, then the titers for cocktail immunization following immunizations with WT become significantly higher. Illustrating this, Figure 6 shows titers as a function of the memory cell fraction given 2 previous WT immunizations. At a memory cell fraction of 0.5, anti-Beta/Gamma titers are around 10 to 15-fold higher for cocktail immunizations than WT immunizations. This is a large difference compared to the memory cell fraction of 1, where the titers are nearly equivalent, suggesting that cocktail immunization is less sensitive to the memory cell fraction. In the WT | WT | cocktail immunization, the WT and cocktail antigens are sufficiently different so that naïve cells can better develop against Beta and Gamma variants than in the WT | WT | WT immunization. Thus, decreasing the memory cell fraction (increasing naïve cell fraction) increases the difference between WT | WT | cocktail immunization and WT | WT | WT immunization. A similar, but less pronounced, effect occurs for anti-Delta titers, as the titers resulting from WT | WT | cocktail immunization are around 1.5 to 2-fold higher than WT | WT | WT immunization. This decrease occurs but is less pronounced for anti-Delta titers because the Delta variant has 2 RBD mutations whereas the Beta and Gamma variants have 3 RBD mutations.

**Figure 6.**
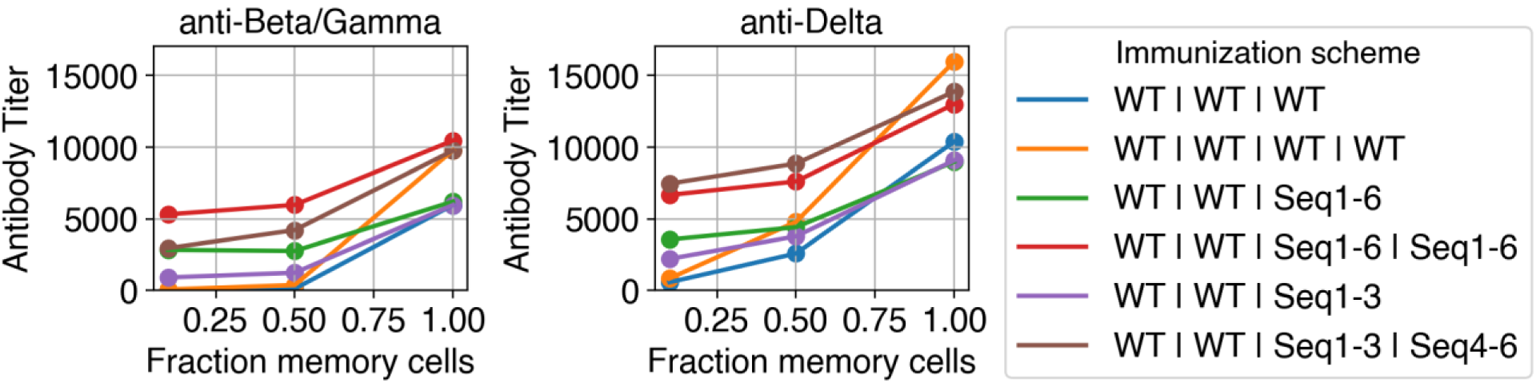
Titers against current variants are affected by the fraction of GC-seeding cells that are memory cells. Titers against Beta/Gamma and Delta variants as a function of the fraction of GC-seeding cells that are memory cells. Beta and Gamma variants carry biochemically similar RBD mutations, so they are equivalent in this model.

### 3.3 The S2’ cleavage site is structurally conserved and is a potential pan-coronavirus vaccine target

Finally, we consider the more general problem of protecting against any coronavirus. To identify a pan-coronavirus vaccine target, we used the conservation analysis outlined in Section 2.1 to calculate spike conservation across 12 coronaviruses from different genera. Figure 7a illustrates conservation using a threshold fraction of 0.8. None of the coronaviruses are conserved in the RBD, which is simply because different coronaviruses bind different receptors. For example, SARS-CoV-2 and SARS-CoV to bind ACE2, while MERS binds DPP4 (Table S1).

**Figure 7.**
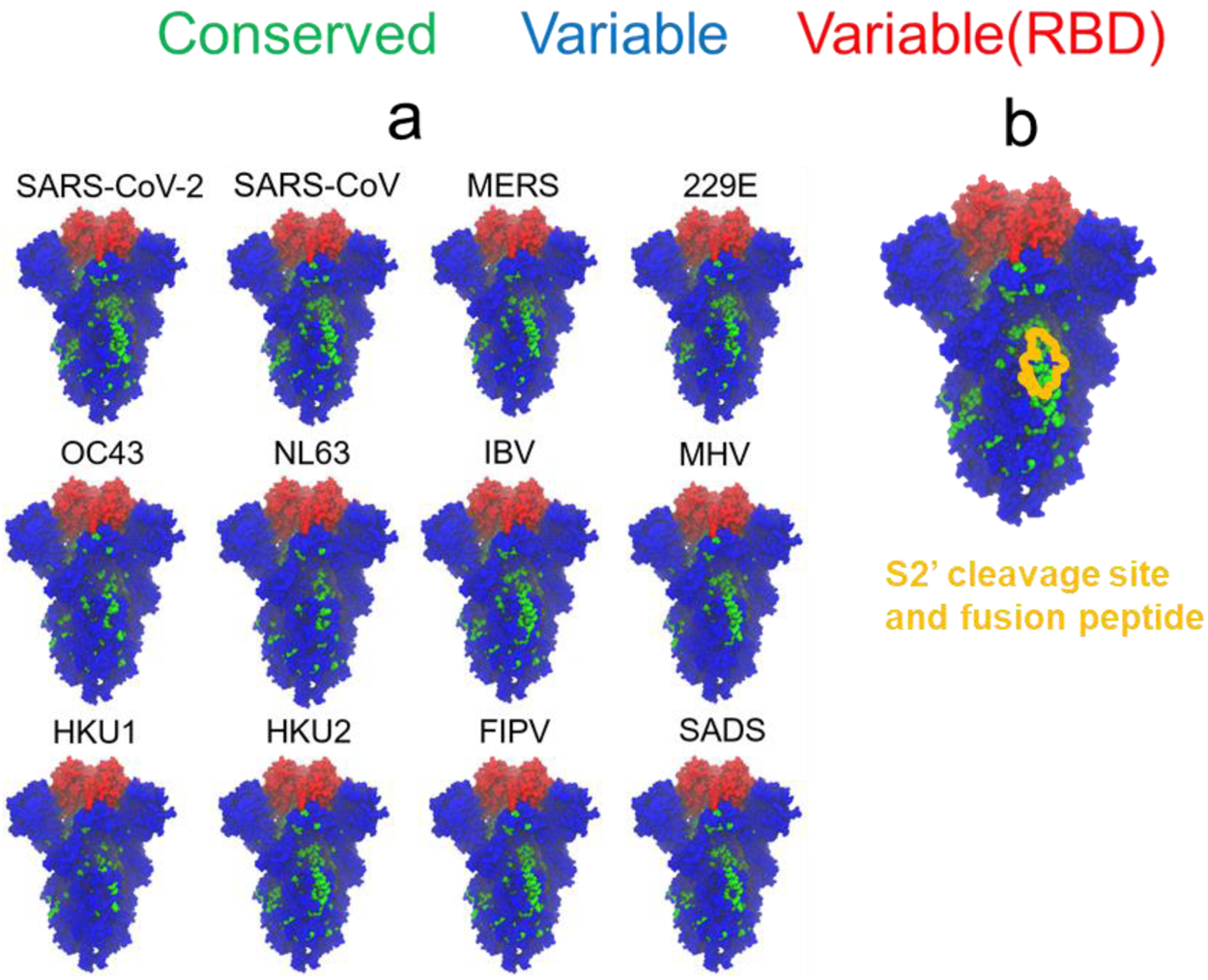
A conserved epitope in the S2 domain overlaps the S2’ cleavage site and fusion peptide. (a) Spike protein structures colored by conservation fraction using each coronavirus as a reference. Green residues have conservation fractions above 0.8, blue residues have conservation fractions below 0.8 and are not in the RBD, and red residues have conservation fractions below 0.8 and are in the RBD. (b) Spike structure colored by conservation fraction using SARS-CoV-2 as a reference. The residues corresponding to the S2’ cleavage site and fusion peptide are outlined in yellow. This structure is shown in a separate panel in order to avoid covering up some residues with the outline.

In contrast to this, the S2 domain of the spike possesses residues that are structurally conserved across many coronaviruses. In particular, 9 out of 12 of the coronaviruses share a conserved patch of residues in the S2 subunit despite coming from diverse lineages. For example, SARS-CoV-2 and IBV both share this conserved region even though SARS-CoV-2 is a betacoronavirus that infects humans and IBV is a gammacoronavirus that infects birds.

These residues are conserved in many coronaviruses because they overlap with the S2’ cleavage site and fusion peptide (Figure 7b). Following S1/S2 cleavage, further cleavage at the S2’ site facilitates the release of the fusion peptide, which fuses the viral and host membranes so that the virus can infect the cell. Since the cleavage site and fusion peptide are functionally important, they must be conserved. It follows that if the virus is subject to evolutionary pressure by S2’-targeting antibodies, it is more difficult to develop escape mutations because potential mutations must also preserve S2’ site function.

Lastly, enzymes such as furin, TMPRSS2, and cathepsin (67–69) must be able to access cleavage sites, which means that the cleavage sites are exposed and accessible to proteins. Furthermore, this limits the virus’s ability to develop spike densities that sterically hinder antibodies from accessing the S2’ site. Electron tomography estimates that SARS-CoV-2 virions have 25-127 spikes and a diameter of 80 nm (70). Even at the dense estimate of 127 spikes, this produces a density of 0.6 spikes per 100 nm^2^, which is half as dense as influenza (1.2 spikes per 100 nm^2^) (25, 71). Thus, compared to the broadly neutralizing epitope on the stem of the influenza hemagglutinin spike, from a steric point of view, generating antibodies to the stem of the coronavirus spike may be more feasible. Taken together, these attributes suggest that targeting the S2’ site could serve as a potential pan-coronavirus vaccine strategy.

## 4 Discussion

SARS-CoV-2 has evolved variants within a few years since its emergence. Current vaccines that immunize with the WT spike show reduced efficacy for some of these variants, especially the Omicron variant. This suggests that vaccines that can protect against multiple variant strains are becoming increasingly important. However, while SARS-CoV-2 is mutable, it is less so than influenza and HIV. Therefore, while universal vaccines against influenza and HIV largely focus on redirecting antibody responses to elicit bnAbs that target conserved regions only, such an approach may not be necessary to protect against SARS-CoV-2 variants. Targeting conserved regions may have disadvantages as well, as antibodies targeting the conserved class 4 RBD epitope can be non-neutralizing (34). In this paper, based on analyses of structures and sequences, we designed antigens that may protect against the most significant antibody-evading variants. Using a neural network, we check that these antigens are viable and can bind ACE2, and so may emerge in the future. We also use a computational model of AM to show that a cocktail of these antigens provides stronger protection against current variants and certain potential variants than the WT vaccination does. In particular, a specific cocktail (sequence 1-6) is predicted to be optimal out of several explored possibilities.

Our cocktail vaccination schemes result in lower titers for the WT antigen than vaccination with the WT antigen alone. It is important to emphasize that our goal is to generate a response that is protective against different variants that may emerge. Such a vaccine will be less protective for any given variant compared to a strain-specific vaccine. However, a vaccine that provides adequate coverage to diverse potential variants would confer protection to a population, which may sufficiently restrict transmission on its own or buy time for a strain-specific vaccine to be developed without the need for strict non-pharmaceutical interventions.

The antibodies elicited by our cocktail vaccine are not bnAbs as they are not designed to target conserved regions. Our AM simulations suggest that the antibodies are broadly protective against multiple mutations that occur in the residues that are mutated in the designed antigens, but not against mutations that occur outside of those residues. Thus, variants that escape our cocktail vaccine may arise. However, note that the residues mutated in the designed antigens are predicted by our conservation analyses to be the most likely to emerge. Also, our AM model weights all residues equally in order to calculate the binding free energy, which is a limitation of the model. In actuality, certain residues easily abrogate antibody binding upon mutation (specifically, class 1 and 2 antibodies elicited from exposure to the WT sequence), and these escape mutations are the ones included in our designed antigens. Thus, our cocktail vaccine should protect against WT escape mutations. The mutations required to evade antibodies generated by our designed vaccine would be complementary to these WT escape mutations. It may be possible that an individual with a previous exposure to the WT sequence could have protection against the mutations that escape our cocktail vaccine.

A recent study created chimeric spike mRNA vaccines, in which the mRNA sequence contains segments from multiple sarbecoviruses, which induced higher neutralizing titers against various sarbecoviruses compared to the WT SARS-CoV-2 sequence (72). However, the chimeras induced lower neutralizing titers against SARS-CoV-2 variants of concern than the WT SARS-CoV-2 sequence. This is not surprising because the chimeras are designed using sarbecoviruses, which are more mutated from the variants of concern than WT SARS-CoV-2 is. If one aims to protect against just SARS-CoV-2 variants, the chimeric vaccine is not better than the WT SARS-CoV-2 vaccine. Our designed antigens are predicted to generate higher titers against SARS-CoV-2 variants than the WT by exploiting data specifically for variants. Therefore, our antigens would be more effective at addressing the immediate threat of vaccine-evading SARS-CoV-2 variants, while chimeric sarbecovirus vaccines would be more effective at addressing the long-term threat of future sarbecoviruses.

Looking beyond sarbecoviruses, a vaccine that protects against any coronavirus would be beneficial. To develop such a pan-coronavirus vaccine, identifying a conserved region of the coronavirus spike to target is necessary. However, previous conservation analysis of coronavirus spikes has not employed available structural data (73), or has not considered diverse sets of coronaviruses (74–76). In this work, we applied our conservation analysis of spike proteins from diverse coronaviruses and found that the S2’ cleavage site and fusion peptide may serve as pan-coronavirus vaccine targets. In support of this idea, several other studies have identified antibodies that bind this epitope (77–80), although not all of these antibodies have potent neutralizing ability. Since the position of the S2’ site is analogous to the position of the stem epitope of influenza hemagglutinin, previous approaches designed to target hemaglutinin’s stem epitope (25, 36, 81–84) can be applied to coronaviruses (85). A limitation of this approach is that the S2’ site is not perfectly conserved, so targeting it would likely yield a vaccine that is less effective against a particular coronavirus than a strain-specific vaccine.

Taken together, the bulk of our results predict that a specific cocktail of variant antigens used as an immunogen would offer protection against variants of SARS-CoV-2 that may emerge in the future. We hope that experimental efforts to test this prediction will follow. We also suggest one vaccination scheme that may help protect against diverse coronaviruses.

## 5 Acknowledgments

Financial support was provided by the National Science Foundation Graduate Research Fellowship Grant No. 1745302, NIH Grant No. 1-R61-AI161805-01, and the Ragon Institute of MGH, MIT, and Harvard. The authors acknowledge the MIT SuperCloud and Lincoln Laboratory Supercomputing Center for providing HPC resources that have contributed to the research results reported within this work.

## 8 Supporting Information Captions

Table S1. Coronaviruses used in conservation analysis with PDB ID, genus, and receptor.

Table S2. Class 1 and 2 antibodies used to identify RBD escape mutations. Text S1. Collection and processing of NCBI coronavirus sequences.

Text S2. Collection and processing of GISAID SARS-CoV-2 sequences. Text S3. Choice of number of escape mutations considered.

Text S4. Titers do not depend on the number of variant antigens encountered in each B cell - FDC interaction

Figure S1. Effect of structural conservation fraction on residue classification as conserved versus variable.

Figure S2. Illustration of the concatenation of individual coronavirus alignments into a single multiple sequence alignment.

Figure S3. Mean panel titers using 10 and 20 seeding cells.

Figure S4. Anti-WT titers for various energy thresholds.

Figure S5. Mean panel titers for 100-antigen and 1000-antigen panels.

Figure S6. GC collapse rate for Seq1-3 | Seq4-6 and Seq1-6 | Seq1-6.

Figure S7. Mean panel titers for different values of mutated residues.

Figure S8. Mean panel titers for different B cell-FDC encounters.

Figure S9. Titers as a function of concentration for different B cell-FDC encounters.

Figure S10. Spike structure colored by conservation fraction exclusively using SARS-CoV-2 data.

