## Supplemental information for "Design of immunogens for eliciting antibody responses that may protect against SARS-CoV-2 variants"

---

***Table S1****. Coronaviruses used in conservation analysis along with the PDB ID of the spike protein, the genus of the coronavirus, and the coronavirus's host receptor.*

| Coronavirus Name | PDB ID | Genus | Receptor |
|---|---|---|---|
| SARS-CoV-2 | 6VXX | Beta | ACE2 (1) |
| SARS-CoV | 5X58 | Beta | ACE2 (2) |
| MERS | 5X59 | Beta | DPP4 (3) |
| 229E | 6U7H | Alpha | APN (4) |
| HKU1 | 5I08 | Beta | Sialic acid (5) |
| OC43 | 6NZK | Beta | Sialic acid (5) |
| NL63 | 7KIP | Alpha | ACE2 (6) |
| IBV | 6CV0 | Gamma | Sialic acid (7) |
| MHV | 3JCL | Beta | CEACAM1 (8) |
| FIPV | 6JX7 | Alpha | APN (9) |
| SADS | 6M16 | Alpha | Unknown |
| HKU2 | 6M15 | Alpha | Unknown |

***Table S2.*** *Class 1 and class 2 antibodies used to identify RBD escape mutations.*

| Antibody Name | Class |
|---|---|
| C105 | 1 (10) |
| COV2-2165 | 1 (11) |
| COV2-2196 | 1 (12) |
| COV2-2832 | 1 (11) |
| LY-CoV016 | 1 (13) |
| REGN10933 | 1 (13) |
| S2E12 | 1 (14) |
| S2H14 | 1 (14) |
| C002 | 2 (10) |
| C121 | 2 (10) |
| C144 | 2 (10) |
| COV2-2050 | 2 (11) |
| COV2-2096 | 2 (11) |
| COV2-2479 | 2 (11) |
| LY-CoV555 | 2 (15) |
| S2D106 | 2 (14) |
| S2H13 | 2 (14) |

| Antibody Name | Class |
| --- | --- |
| S2H58 | 2 (14) |
| S2X16 | 2 (14) |
| S2X58 | 2 (14) |

### S1. Collection and processing of NCBI coronavirus sequences

Coronavirus spike protein sequences were obtained by searching for the coronavirus name in the NCBI Protein database and downloading spike protein sequences. Sequences uploaded as of May 20, 2021 were obtained as this was when the download was done. Then, sequences not corresponding to the spike protein of the appropriate coronavirus were manually removed based on the organism and protein name, leading to ~4000 coronavirus sequences in total. The sequences of a particular coronavirus were aligned to the sequence of the spike protein PDB structure (PDB structure code is in Table S1) using the ClustalW algorithm (16) in MEGAX (17) with gap opening penalty of 10 and gap extension penalty of 0.2 (sequences available at https://github.com/ericzwang/sars2-vaccine/blob/main/data/aligned-cov-sequences.gz).

### S2. Collection and processing of GISAID SARS-CoV-2 sequences

SARS-CoV-2 genomes with collection dates between December 30, 2019 and July 27, 2021 were downloaded from the GISAID database (~3.3M total). Each genome was translated into its amino acid sequence in all three reading frames using Biopython version 1.78 (18). The proper reading frame for each genome was considered to be the one that contains the subsequence MFVFLVLLP, the first 9 amino acids of the spike protein. If none of the reading frames contained this subsequence, then the genome was discarded. The end of the spike protein was found by identifying the next stop codon after the subsequence. Using the beginning and end of the spike protein thus identified, the sequence of the spike protein was extracted. As a quality check, spike protein sequences were discarded if they were not between 1270 and 1273 amino acids long, or if they contained 3 or more ambiguous amino acids, leading to ~300,000 high quality sequences. These sequences were then aligned with the Wuhan reference sequence (NCBI: NC_045512.2) using Clustal Omega version 1.2.4 (19) with default settings (Clustal Omega was used instead of ClustalW because the number of sequences was very large), and alignment gaps in the Wuhan reference sequence were removed to produce the final alignment. Sequences are not provided due to GISAID terms of service.

### S3. Choice of number of escape mutations considered

We chose the top 34 escape mutations because mutating too many residues was likely to destabilize the RBD. First, we noted that the deep mutational scanning data was based on RBDs with 1-7 mutations, so we could not design RBDs with more than 7 mutations. Furthermore, rather than mutating the maximum number of 7 residues, we chose to mutate 5 residues per RBD to further decrease the probability of destabilizing the RBD. Since the mutations were divided into 2 groups, mutating 5 residues per RBD allowed 10 unique residues to be mutated in total. The top 34 escape mutations was the largest set that contained 10 unique residues (the top 35 escape mutations contained 11 unique residues).

If 7 mutations per RBD were allowed, this corresponded to 14 unique residues. The top 40 escape mutations contained 14 unique residues. However, the total escape of the top 40 escape mutations was only moderately larger than the total escape of the top 34 escape mutations, so choosing the top 34 mutations did not significantly affect the escape potential of our antigens.

### S4. Titers do not depend on the number of variant antigens encountered in each B cell - FDC interaction

We also simulated the case in which antigens are heterogeneously distributed on the FDC, so a B cell encounters only one randomly chosen antigen in each cycle. In Figure S9, the mean panel titers are not significantly different from the all-antigen case. The main difference between the all-antigen and one-antigen case is that the optimal concentration is lower for the all-antigen case (Figure S10). An optimal concentration exists because low concentrations induce GC collapse while high concentrations are incapable of discriminating between low-affinity and high-affinity clones.

In the one-antigen case, a B cell that has high affinity to a particular antigen will likely survive a cycle if it encounters that antigen on the FDC, but it will likely die if it encounters a different antigen for which the affinity is below a threshold. Over many cycles, the B cell is likely to encounter a different antigen at some point and die. Increasing the concentration counteracts this because a higher concentration will decrease the probability of low-affinity B cells dying. Thus, the optimal concentration is higher for the one-antigen case. It is important to note that our goal in this study is not to design antigens and immunization schemes that result in bnAbs that focus their binding footprint only on the conserved residues and can protect against a large set of variants. Having a high concentration of antigen in the one-antigen case would result in a polyclonal response of strain-specific antibodies, just like the all-antigen case, not bnAbs. Our goal in this study is to elicit a polyclonal antibody response that can protect against a limited set of variants. For this situation, manipulating antigen concentration can mitigate the difference between heterogeneous and homogeneous distribution of antigens on FDCs.

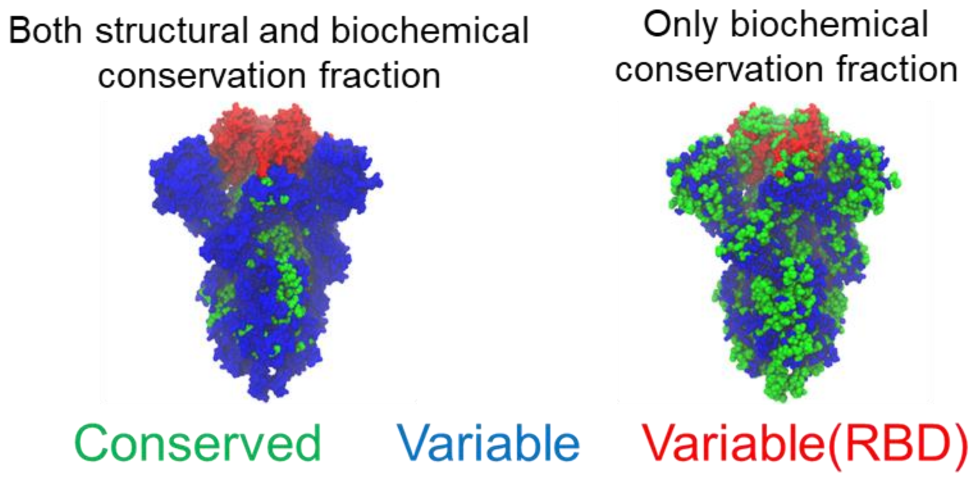


***Figure S1***. *Spike protein structures colored by conservation fraction (either as an average of the structural and biochemical conservation fraction or only the biochemical conservation fraction) using SARS-CoV-2 as a reference. Green residues have conservation fractions above 0.8, blue residues have conservation fractions below 0.8 and are not in the RBD, and red residues have conservation fractions below 0.8 and are in the RBD.*

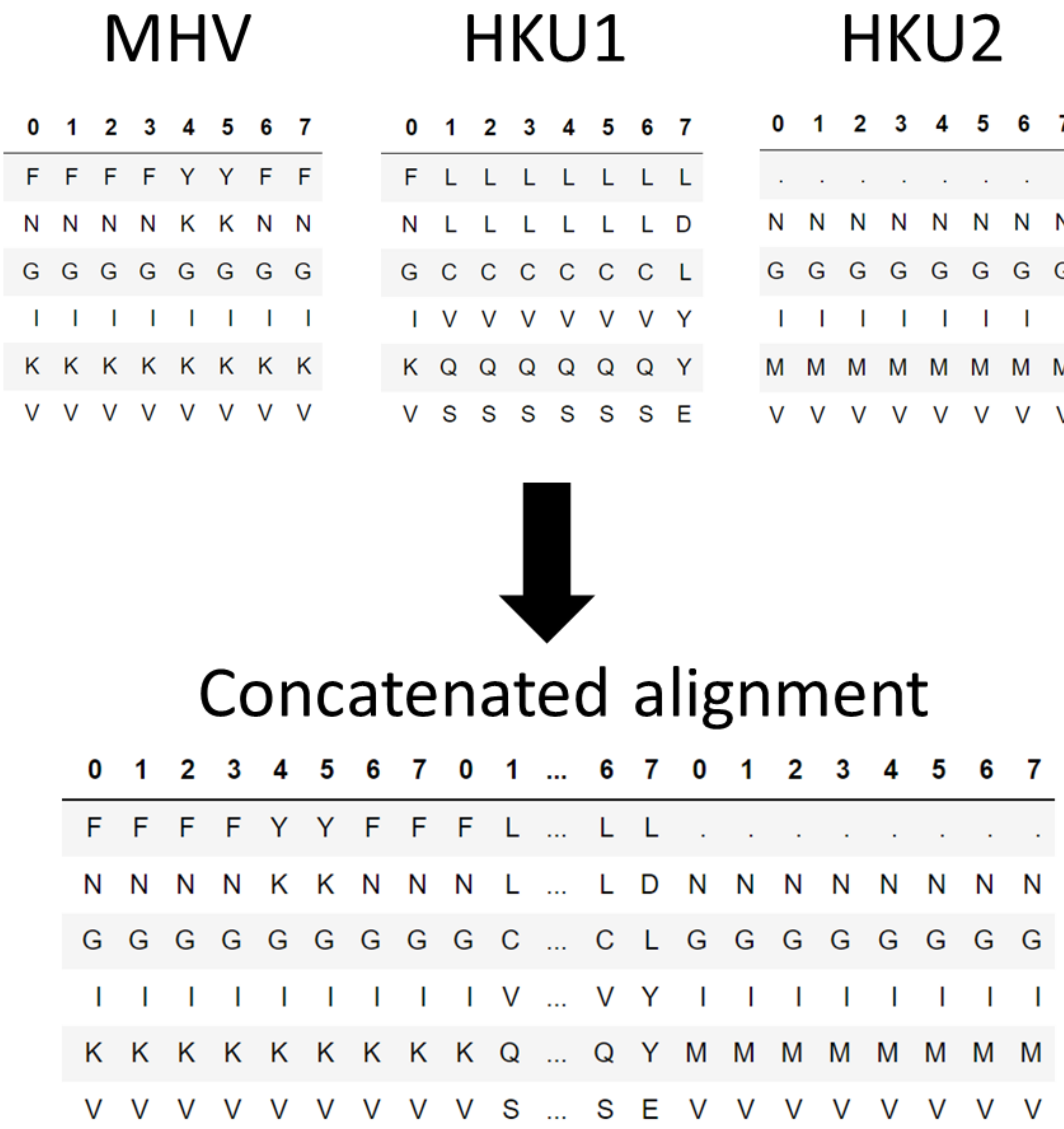


***Figure S2**. Illustration of the concatenation of individual coronavirus alignments into a single multiple sequence alignment. The number of coronaviruses and size of the alignments have been reduced for clarity and to match the example table from Figure 1.*

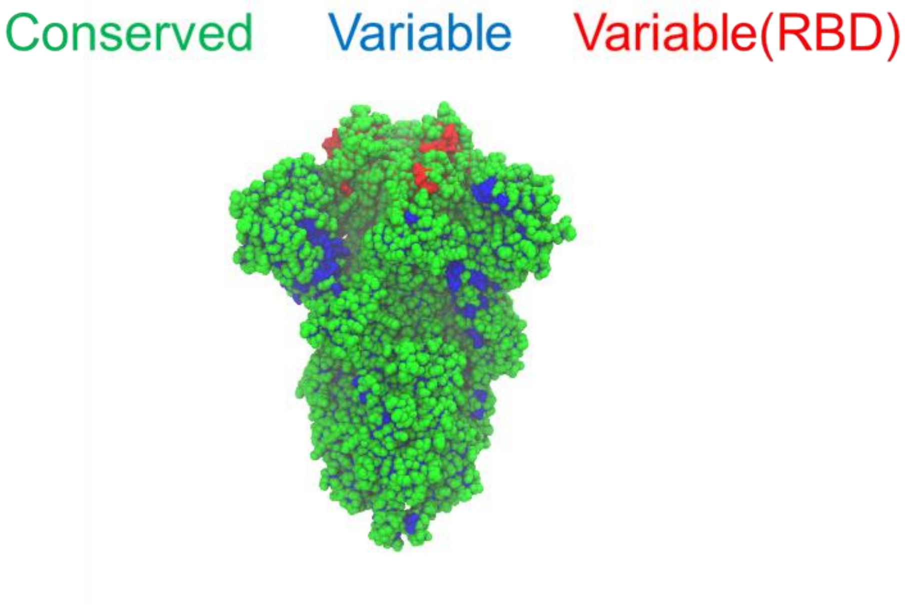


***Figure S3.*** *Spike structure colored by conservation fraction exclusively using SARS-CoV-2 data. Green residues have conservation fractions above 0.8, blue residues have conservation fractions below 0.8 and are not in the RBD, and red residues have conservation fractions below 0.8 and are in the RBD. PDB structures used to calculate the structural conservation fraction include the WT structure (PDB ID: 6VXX), the Alpha variant structure (PDB ID: 7LWI), the Beta variant structure (PDB ID: 7LWS), the Delta variant structure (PDB ID: 7V7Q), and the Gamma variant structure (PDB ID: 7M8K). ~300,000 spike sequences (obtained from GISAID as described above) were used to calculate the biochemical conservation fraction.*

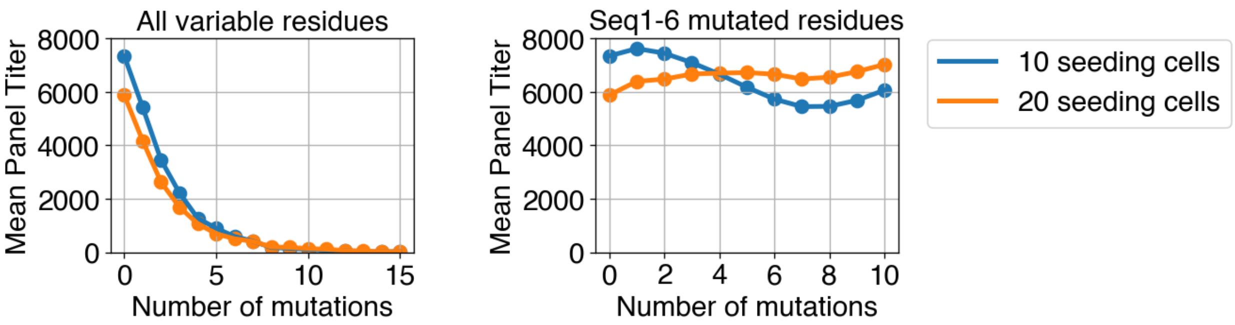


***Figure S4.*** *Mean panel titers for the Seq1-6 | Seq1-6 immunization scheme using 10 and 20 seeding cells. Mutations occur either in any variable residues (All variable residues) or in the same residues that are mutated in the sequences 1-6 (Seq1-6 mutated residues). B cells are assumed to encounter all antigens at a time on the FDC.*

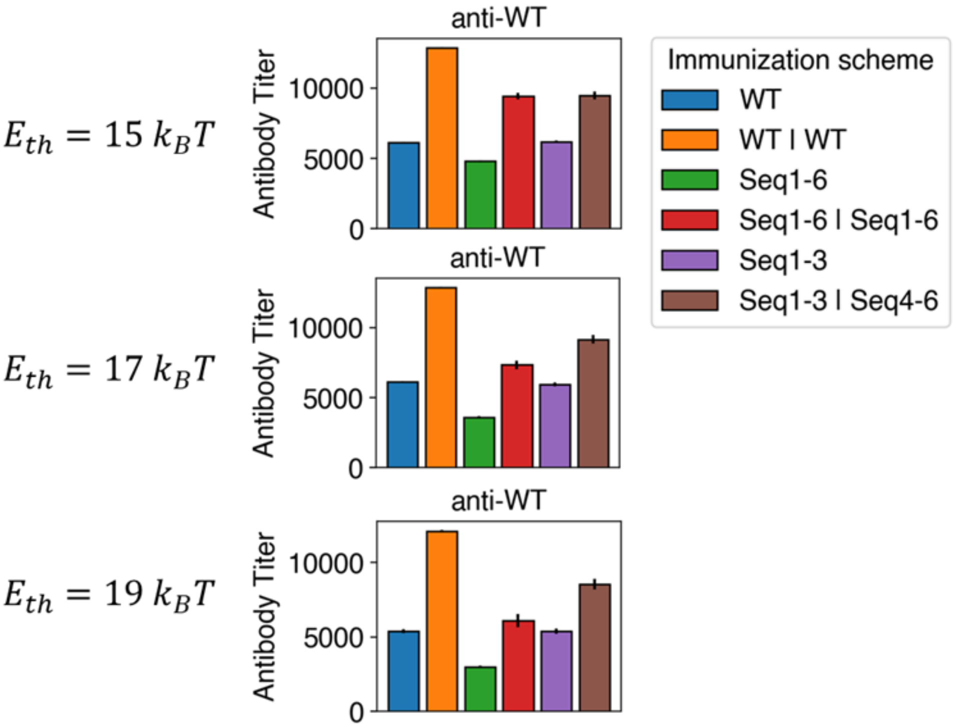


***Figure S5.*** *Anti-WT titers for various immunization schemes and energy thresholds for calculating titers ($E_{th}$).*

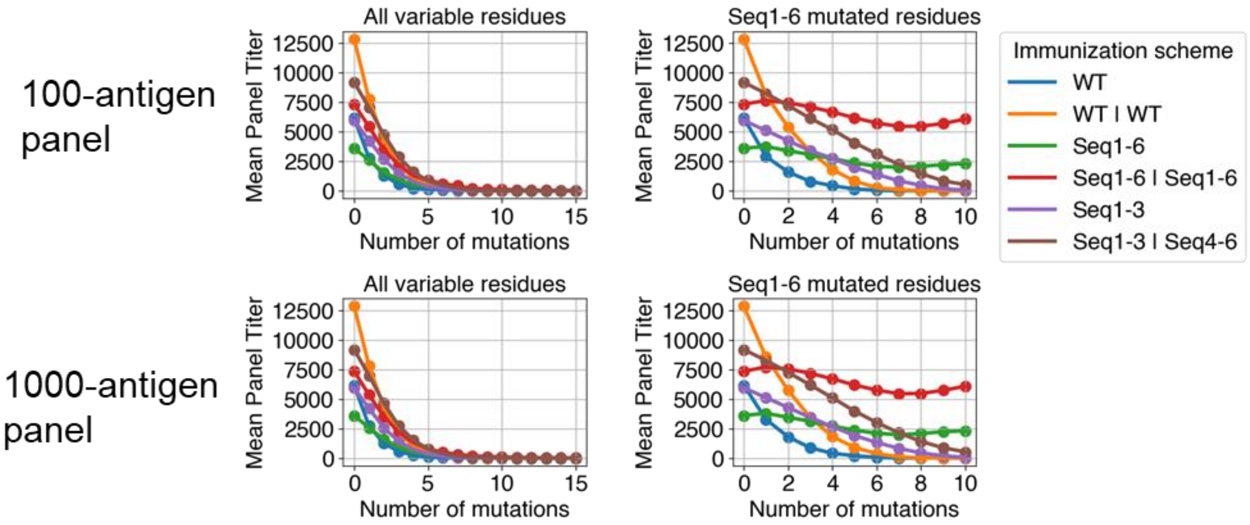


***Figure S6.*** *Mean panel titers as a function of the number of mutations in panel antigens. Mutations occur either in any variable residues (All variable residues) or in the same residues that are mutated in the sequences 1-6 (Seq1-6 mutated residues). Panel titers are calculated against panels of 100 antigens and 1000 antigens. Mutated residues take on a value of -4. B cells are assumed to encounter all antigens at a time on the FDC.*

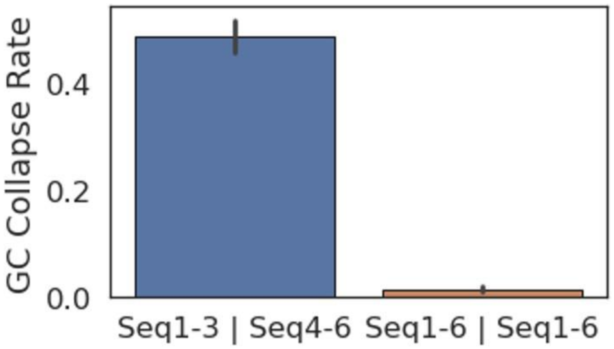


***Figure S7.*** *GC collapse rate for Seq1-3* | *Seq4-6 and Seq1-6* | *Seq1-6*

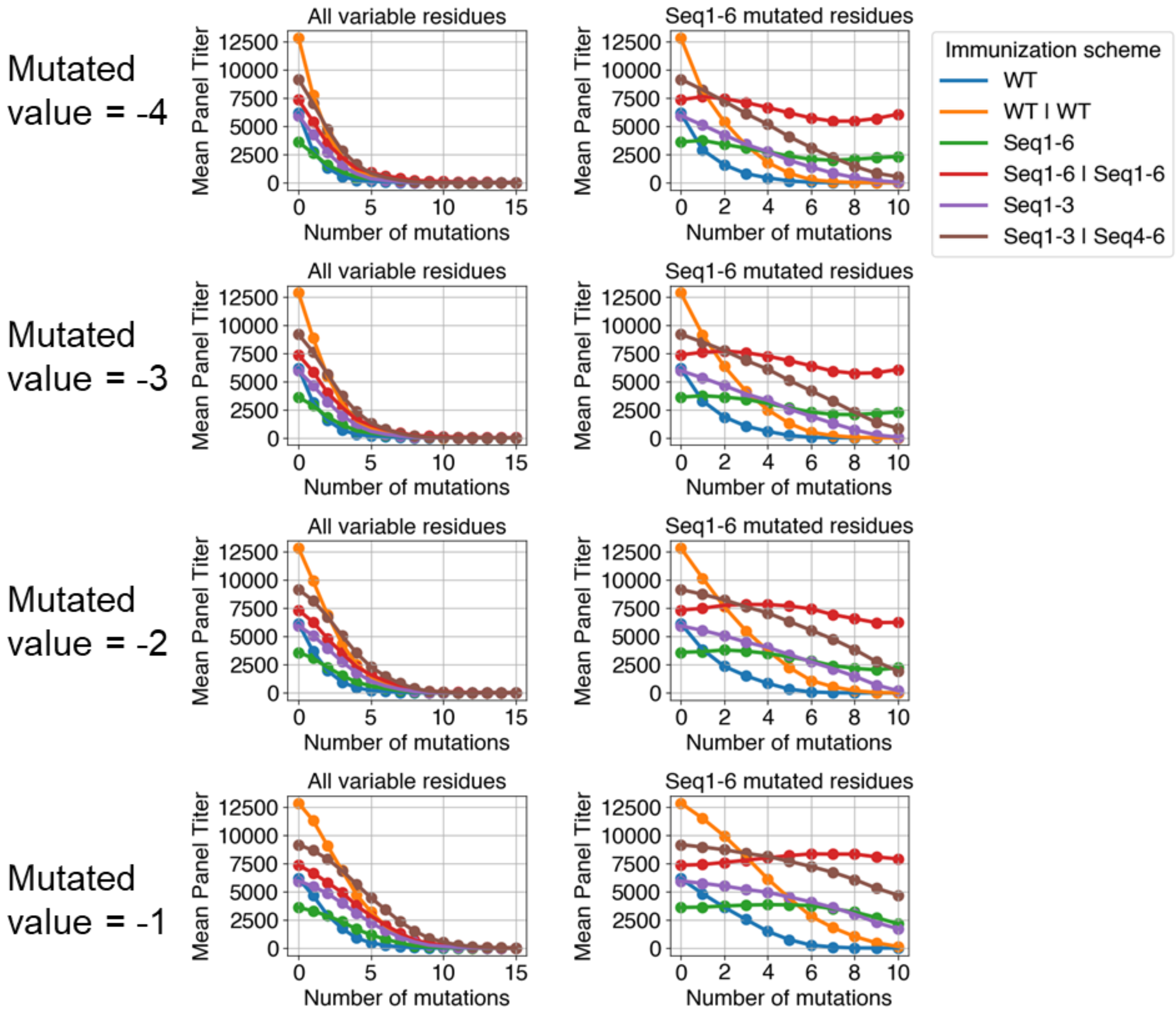


***Figure S8.*** *Mean panel titers as a function of the number of mutations in panel antigens. Mutations occur either in any variable residues (All variable residues) or in the same residues that are mutated in the sequences 1-6 (Seq1-6 mutated residues). B cells are assumed to encounter all antigens at a time on the FDC. Mutated residues take on values of -4, -3, -2, or -1, as indicated.*

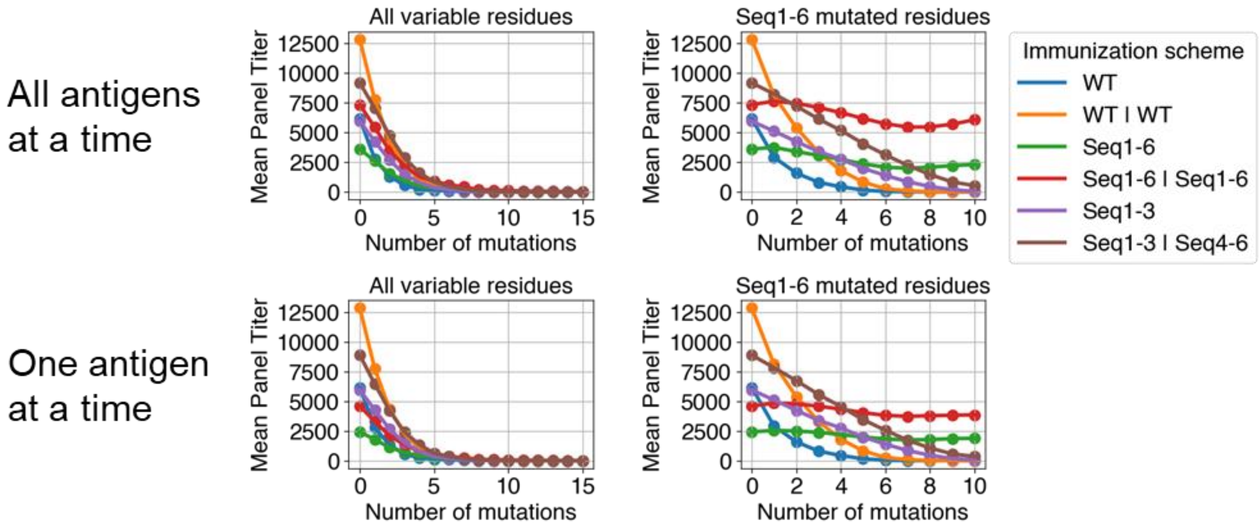


***Figure S9.*** *Mean panel titers as a function of the number of mutations in panel antigens. Mutations occur either in any variable residues (All variable residues) or in the same residues that are mutated in the sequences 1-6 (Seq1-6 mutated residues). Mutated residues take on values of -4. B cells encounter either all antigens at a time (All-antigen) or one antigen at a time (One-antigen) on the FDC.*

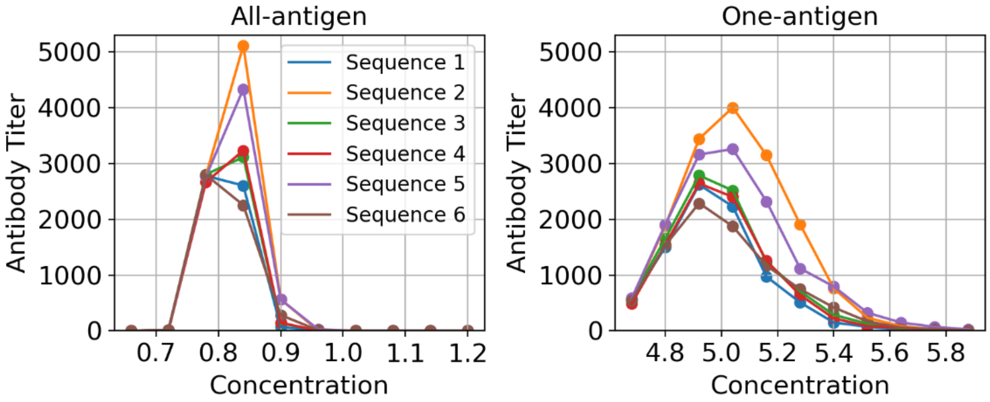


***Figure S10.*** *Antibody titers against sequences 1-6 as a function of concentration if B cells encounter all antigens (All-antigen) or one antigen (One-antigen) at a time on the FDC.*
